# Exposure of Greenlandic Inuit and South African VhaVenda men to the persistent DDT metabolite is associated with an altered sperm epigenome at regions implicated in paternal epigenetic transmission and developmental disease – a cross-sectional study

**DOI:** 10.1101/2022.08.15.504029

**Authors:** A. Lismer, X. Shao, M.C. Dumargne, C. Lafleur, R. Lambrot, D. Chan, G. Toft, J.P. Bonde, A.J. MacFarlane, R. Bornman, N. Aneck-Hahn, S. Patrick, J.M. Bailey, C. de Jager, V. Dumeaux, J.M. Trasler, S. Kimmins

**Affiliations:** Department of Pharmacology and Therapeutics, Faculty of Medicine and Health Sciences, McGill University, Montreal, QC H3G 1Y6, Canada; Digital Technologies Research Centre, National Research Council Canada, Ottawa, ON K1A 0R6, Canada; Department of Animal Science, Faculty of Agricultural and Environmental Sciences, McGill University, Montreal, QC H9X 3V9, Canada; Child Health and Human Development Program, Research Institute of the McGill University Health Centre, Montreal, QC H4A 3J1, Canada; Steno Diabetes Center Aarhus, Aarhus University Hospital, Aarhus, DK-8200 Aarhus N, Denmark; Department of Occupational and Environmental Medicine, Bispebjerg University Hospital, Copenhagen, DK-2400 Copenhagen NV, Denmark; Institute of Public Health, University of Copenhagen, Copenhagen, DK-1016 Copenhagen K, Denmark; Nutrition Research Division, Health Canada, Ottawa, ON K1A 0K9, Canada; Environmental Chemical Pollution and Health Research Unit, Faculty of Health Sciences, School of Health Systems and Public Health, University of Pretoria, Pretoria, 0084, South Africa; University of Pretoria Institute for Sustainable Malaria Control, School of Health Systems and Public Health, Faculty of Health Sciences, University of Pretoria, 0028, South Africa; Research Centre on Reproduction and Intergenerational Health, Department of Animal Sciences, Université Laval, Quebec City, QC G1V 0A6, Canada; Departments of Anatomy & Cell Biology and Oncology, Western University, London, Ontario, N6A 3K7, Canada; Department of Human Genetics, Faculty of Medicine, McGill University, Montreal, QC H3A 0C7, Canada; Department of Pediatrics, Faculty of Medicine, McGill University, Montreal, QC H4A 3J1, Canada

**Author notes:** These authors contributed equally to this work (X. Shao, M.C. Dumargne).

## Abstract

**Background:** The persistent organochlorine dichlorodiphenyltrichloroethane (DDT) is banned world-wide due to its negative health effects and persistence in the environment. It is exceptionally used as an insecticide for malaria control. Exposure occurs in regions where DDT is applied, as well as in the arctic where it’s endocrine disrupting metabolite, *p,p’-*dichlorodiphenyldichloroethylene (*p,p’-*DDE) accumulates in marine mammals and fish. DDT and *p,p’-*DDE exposures are linked to birth defects, infertility, cancer, and neurodevelopmental delays. Of particular concern is the potential of DDT use to impact the health of generations to come. Generational effects of toxicant exposures have been described in animal models and implicated germline epigenetic factors. Similar generational effects have been shown in epidemiological studies. Although advances in understanding the molecular mechanisms mediating this epigenetic inheritance have been made, there remain major knowledge gaps in how this occurs in humans. In animal and human models, DNA methylation (DNAme) has been implicated in paternal epigenetic effects. In animal models, histone H3K4 trimethylation (H3K4me3) has been shown to be responsive to the paternal environment and linked with epigenetic transmission to the embryo. Our objectives were to define the associations between *p,p’-*DDE serum levels and alterations in the sperm methylome and H3K4me3 enrichment using next generation sequencing. We aimed to compare regions of epigenomic sensitivity between geographically diverse populations with different routes and levels of exposures, and to identify interactions between altered DNAme and H3K4me3 regions. The potential for *p,p’-*DDE to impact the health of the next generation was explored by examining the functions of the genomic regions impacted, their roles during embryo development, and in health and disease.

**Methods:** In the Limpopo Province of South Africa, we recruited 247 VhaVenda South African men from 12 villages that either used indoor residual spraying with DDT for malaria control or not. We selected 49 paired blood and semen samples, from men that ranged from 18 to 32 years of age (mean 25 years). Sample inclusion was based on normal sperm counts (> 15 million/ml), normal sperm DNA fragmentation index, and testing a range of *p,p’-*DDE exposure levels (mean 10,462.228 ± 1,792.298 ng/ml). From a total of 193 samples, 47 Greenlandic Inuit blood and semen paired samples were selected from the biobank of the INUENDO cohort. The subjects ranged from 20 to 44 years of age (mean 31 years), were born in Greenland, and all had proven fertility. Sample selection was based on obtaining a range of *p,p’*-DDE exposure levels (mean 870.734 ± 134.030 ng/ml). Here we determined the molecular responses at the level of the sperm epigenome to serum *p,p’*-DDE levels using MethylC-Capture-seq (MCC-seq) and chromatin-immunoprecipitation followed by sequencing (ChIP-seq). We identified genomic regions with altered DNA methylation (DNAme) and differential enrichment of histone H3 lysine 4 trimethylation (H3K4me3) in sperm. We used *in silico* analyses to discover regions of differential methylation associated with *p,p’-*DDE levels that were predicted to be transmitted and persist in the embryo.

**Results:** Alterations in DNAme and H3K4me3 enrichment followed dose response-like trends, and we identified overlapping genomic regions with DNAme sensitivities in both populations. Altered DNAme and H3K4me3 in sperm occurred at transposable elements and regulatory regions involved in fertility, disease, development, and neurofunction. A subset of regions with altered sperm DNAme and H3K4me3 were predicted to persist in the pre-implantation embryo and were associated with embryonic gene expression.

**Limitations:** The samples were collected from remote areas of the world thus sample size is relatively small. The populations differed in the routes of exposure, timing of collection, mean age (mean of 25 versus 31 years of age in South African and Greenlandic populations respectively) and in the timing of *p,p’-*DDE measurement. Moreover, the Greenlandic Inuit men were proven fertile whereas the fertility status of the South African men was unknown. Confounding factors such as other environmental exposures and selection bias cannot be ruled out.

**Conclusions:** These findings suggest that in men, DDT and *p,p’-*DDE exposure impacts the sperm epigenome in a dose-responsive manner and may negatively impact the health of future generations through epigenetic mechanisms.

## Introduction

Dichlorodiphenyltrichloroethane (DDT) is a lipophilic organochlorine pollutant that is listed under the Stockholm Convention, with the objective to eliminate its use to protect human health and the environment. However, its use persists in Africa and India. Consequently, DDT continues to damage health and accumulates in the environment, becoming particularly concentrated in the aquatic food chain (Turusov et al., 2002; van den Berg et al., 2017). Currently, a sanctioned use of DDT is for indoor residual spraying to control malarial disease-vectors *Anopheles gambiae* and *Anopheles arabiensis* (Himeidan et al., 2007; World Health Organization. Global Malaria and World Health Organization. Malaria, 2006). DDT is a potent endocrine disruptor with DDT isomers exerting different endocrine disrupting actions: *p,p’-*dichlorodiphenyldichloroethylene (*p,p’-*DDE) is antiandrogenic whereas *p,p’*-DDT and *o,p’*-DDT are estrogenic (Bitman et al., 1968; Kelce et al., 1995). The main DDT metabolite, *p,p’-*DDE is associated with adverse health outcomes, such as low birth weight, neurotoxicity, cancer, poor semen quality, and increased risk of preterm birth (Aneck-Hahn et al., 2007; Bornman et al., 2010; Bouwman and Kylin, 2009; Cartier et al., 2014; Eskenazi et al., 2009; Longnecker et al., 2001). Animal models and epidemiological studies indicate that paternal exposures to organochlorines and DDT, can impact health intergenerationally and potentially transgenerationally (Skinner et al., 2018; Van Oostdam et al., 2005; Weihe et al., 2016). These generational effects have been attributed to the epigenome, which refers to the biochemical content associated with DNA that impacts chromatin organization as well as gene expression and is transmitted via the gametes to alter phenotypes across generations (Lismer et al., 2021; Lismer et al., 2020; Siklenka et al., 2015). The epigenome includes: 1) DNA methylation (DNAme) occurring at the 5′-position of cytosine residues within CpG dinucleotides; 2) the modifications on nucleosome proteins, histones, such as methylation, acetylation, and phosphorylation among others; and 3) small non-coding RNAs. A plausible molecular mechanism for paternal epigenetic inheritance is sperm-mediated transmission of regions bearing environmentally altered DNA and histone methylation.

In mammals, spermatogenesis entails unique testis-specific gene expression programs that are accompanied by dynamic remodeling of the chromatin (Kimmins and Sassone-Corsi, 2005; Lambrot et al., 2019; Larose et al., 2019; Maezawa et al., 2018). During this process, the majority of histones are replaced by protamines, with 1% of histones retained in sperm from mice and 15% in men (Erkek et al., 2013; Hammoud et al., 2009; Kimmins and Sassone-Corsi, 2005). Retained histones are conserved across mammalian species and are found at gene regulatory regions implicated in spermatogenesis, sperm function, embryo development, metabolism and routine cellular processes (Brykczynska et al., 2010; Hammoud *et al*., 2009; Lesch et al., 2016). We have shown in human and mouse sperm that the gene activating modification, histone H3 lysine 4 trimethylation (H3K4me3) localizes to genes involved in fertility, metabolism and development (Lambrot et al., 2021; Lismer *et al*., 2021; Lismer *et al*., 2020; Pepin et al., 2022). During spermatogenesis, disrupting the function of histone modifiers genetically or via environmental exposures, as well as modifying histone residues in the paternal pronucleus of the zygote, provides compelling evidence that sperm-transmitted histones serve critical functions in embryo development and offspring health (Aoshima et al., 2015; Lesch et al., 2019; Lismer *et al*., 2021; Lismer *et al*., 2020; Pepin *et al*., 2022; Santenard et al., 2010; Siklenka *et al*., 2015; Stringer et al., 2018). A clear indicator that sperm histones are transmitted at fertilization and retained in the embryo is the localization of sperm-enriched histone variant H3.3 to the paternal pronucleus of the zygote (Aoshima *et al*., 2015; Ishiuchi et al., 2021; Santenard *et al*., 2010). Whether histones in sperm from men, specifically histone H3K3me3, are similarly transmitted to alter embryonic gene expression is unknown. In support of this possibility, H3K4me3 peaks in human sperm occur at genes that are expressed in the pre-implantation embryo (Lambrot *et al*., 2021).

Human sperm DNA is highly methylated (Molaro et al., 2011) and disruption of methyl signatures is related to infertility and abnormal offspring development (Bestor, 1998; Bourc’his and Bestor, 2004; Okano et al., 1999). Promoters with high CpG content are enriched for nucleosomes and predominantly hypomethylated in sperm (Erkek *et al*., 2013; Hammoud *et al*., 2009). Although sperm epigenetic modifications that respond to environmental exposures are most frequently studied alone, there are likely interdependent interactions between DNA methylation and chromatin in sperm. For example, there is cooperativity between chromatin and DNA methylation machinery; DNMT3L acts via the amino terminus of H3K4me to recruit the *de novo* DNA methyltransferase. When H3K4 is methylated, this interaction with DNMT3L is abrogated and DNA methylation is blocked (Ooi et al., 2007). Whole-genome bisulfite sequencing and H3K4me3 ChIP-seq of human sperm revealed that over 24,000 H3K4me3 peaks coincided regions of intermediate (between 20 and 80%) and high (over 80%) DNA methylation (Lambrot *et al*., 2021). These overlapping regions occurred at genes implicated in fertility and development (Lambrot *et al*., 2021). How sperm DNAme and chromatin respond in concert to environmental exposures including endocrine disrupting chemicals (EDCs) remains underexplored. It is also unknown whether common H3K4me3 and DNAme regions are sensitive to DDT and *p,p’-* DDE, and could be transmitted and retained in the embryo.

The use of animal models in combination with advances in epigenomic techniques have directly implicated errors in the establishment and maintenance of the germline epigenome to birth defects, neurodevelopmental disorders, and common diseases such as diabetes and cancer (Harutyunyan et al., 2019; Lismer *et al*., 2021; Lismer *et al*., 2020; Michealraj et al., 2020; Pepin et al., 2021; Siklenka *et al*., 2015). In mice, sperm DNAme and histone modifications can be altered by environmental exposure to toxicants such as those found in insecticides and plastics, but also by obesity and nutrient restriction (Donkin et al., 2016; Lambrot et al., 2013; Lismer *et al*., 2021; Maurice et al., 2021; Radford et al., 2014). These epigenetic errors in sperm can subsequently be transmitted to the embryo to alter embryonic gene expression, development, and offspring health (Lismer *et al*., 2021; Radford *et al*., 2014). In rodents, exposure to organochlorines including DDT, has been linked with altered DNAme in sperm and associated with poor reproductive outcomes, and offspring metabolic disease (Herst et al., 2019; King et al., 2019; Lessard et al., 2019). In a prior study using the same Greenlandic Inuit population than here, *p,p’-*DDE exposures were examined in relation to sperm DNA methylation by targeted pyrosequencing at Satα tandem repeat sequences, and at LINE-1 and Alu transposable elements. Global DNA methylation levels were also measured by flow cytometric fluorescence. Using these approaches, the results were inconclusive as to whether sperm DNAme is altered by *p,p’-*DDE exposures (Consales et al., 2016). Therefore, whether the sperm epigenome of men is impacted by DDT and *p,p’-*DDE exposures and could be associated with paternal epigenetic transmission remains unresolved.

Given the improvement in epigenome-wide sequencing approaches for epigenome-wide association studies, we revisited the response of the sperm methylome to DDT and *p,p’-*DDE. In this study we aimed to identify alterations in sperm DNAme and histone H3K4me3 associated with levels of the persistent DDT metabolite measured in blood and how these epigenetic alterations could be implicated in epigenetic inheritance. We studied two geographically diverse exposed populations, Greenlandic Inuit and South African VhaVenda men. To quantitatively identify regions that were altered in DNAme and histone methylation of exposed men, we used a sperm customized methyl-capture approach followed by sequencing (MCC-seq; Greenlandic and South African populations), and chromatin immunoprecipitation targeting histone H3K4me3 followed by sequencing (ChIP-seq; South African population exclusively). We then performed differential and functional analyses to define altered DNAme and H3K4me3 regions associated with high levels of *p,p’-*DDE exposure. *In silico* analyses were used to further explore the possibility that these epigenetic alterations in sperm could be transmitted to the embryo at fertilization, impact embryonic gene expression, and persist throughout embryonic development.

## Materials and Methods

### Ethical statements

For the Greenlandic Inuit cohort of the INUENDO biobank, the Scientific Ethics Committee for Greenland approved the research protocol (project reference number 2014-25/26). For the South African VhaVenda cohort, the research protocol was approved by the Scientific Ethics Committee of the Faculty of Health Sciences, University of Pretoria and the Limpopo Provincial Government’s Department of Health approved the research protocol (project reference number 43/2003), by Health Canada and Public Health Agency of Canada’s Research Ethics Board’s (project reference number REB 2019-0006) and the Ethics Committee for the Faculty of Medicine and Health Sciences, McGill University (project reference number A09-M57-15B).

### Greenlandic Inuit population blood and semen collection

Indigenous Arctic inhabitants consume a traditional marine mammal diet consisting of whale, walrus and seal, and are at a high risk of exposure to *p,p’-*DDE through its bioaccumulation in the marine food chains (Brown et al., 2018; Muir et al., 1992). Greenlandic Inuit blood and semen paired samples were selected from the biobank of the INUENDO cohort (Spanò et al., 2005) The subjects ranged from 20 to 44 years of age (mean age of 31 years), were born in Greenland and all had proven fertility with confirmation of a pregnant partner. Sample selection was based on obtaining a range in *p,p’-*DDE exposure levels (mean 870.734 ± 134.030 ng/ml, Fig. S1 and Table S1A) (n = 47 from 193 total for MCC-seq). Full details on recruitment and the cohort have been previously described (Spanò *et al*., 2005). Data was available on smoking, DNA fragmentation index and Body Mass Index (BMI) (see aggregate data Table S1A). Note for adherence with General Data Protection Regulations (GDPR) individual data cannot be published. Semen samples from participants who gave informed consent were collected between May 2002 and February 2004 by masturbation in private room. The men were asked to abstain from sexual activities for over 2 days before collecting the sample. Immediately after collection, semen samples were kept close to the body to maintain a 37°C temperature when transported to the laboratory. Two cryotubes with 0.2 ml aliquots of undiluted raw semen collected 30 min after liquefaction, were prepared from each semen sample, and long-term storage was at −80°C freezer. Venous blood samples were collected up to 1 year in advance of the semen collection. The blood samples were centrifuged immediately after collection and sera were stored in a −80°C freezer for later analysis. Samples were analyzed at the department of Occupational and Environmental Medicine in Lund, Sweden as previously described (Jönsson et al., 2005; Richthoff et al., 2003) (Table S1A). Briefly, the sera were transported on dry ice to the Department of Occupational and Environmental Medicine in Lund, Sweden, where all the analyses of *p,p’-*DDE were performed (see aggregate data Table S1A). The *p,p’-*DDE was extracted by solid phase extraction using on-column degradation of the lipids and analysis by gas chromatography mass spectrometry as previously described (Rignell-Hydbom et al., 2004). The relative standard deviations, calculated from samples analyzed in duplicate at different days, were 1% at 1 ng/mL (n = 1,058), 8% at 3 ng/mL (n = 1,058) and 7% at 8 ng/mL (n = 1,058) and the detection limit was 0.1 ng/mL for *p,p’-*DDE.

### South African population blood and semen collection

The Vhembe district is a malaria endemic area where housing includes mud, or brick or cement dwellings that are sprayed with DDT, or not to control for malaria. Participants volunteered from 12 villages from the Vhembe district of the Limpopo province of South Africa that were either sprayed (n = 32) or not non-sprayed (n = 17), This prospective study was conducted in the same manner as our prior studies and full details on recruitment methods and the questionnaire have been previously described (De Jager et al., 2009). Sample collection occurred in October 2016, February 2017, and November 2017. Men were excluded from the study if they were less than 18 or more than 40 years of age, appeared intoxicated, or had a neuropsychiatric illness. Physical measurements included height and weight. All participants provided informed consent and were interviewed using a questionnaire on their use of insecticides, diet, smoking, drinking, and drug use (see aggregate data Table S1B). Fertility status was not queried. Based on a cross-section of 247 men, we selected 49 paired blood and semen samples, from men that ranged from 18 to 32 years of age (mean 25 years). Sample inclusion was based on normal sperm counts (> 15 million/ml), normal sperm DNA fragmentation index, and testing a range of *p,p’-*DDE exposure levels (mean 10,462.228 ± 1,792.298 ng/ml, Fig. S1 and Table S1B). Note for adherence with General Data Protection Regulations (GDPR) individual data cannot be published. The semen was preserved in Sperm Freeze (LifeGlobal) and stored in liquid nitrogen for transport, followed by long-term storage in a −80°C freezer. Semen analysis was performed according to the World Health Organization standard (Organisation, 1999) and the DNA fragmentation index was determined (Table S1B). Serum lipid measurements and analysis for *p,p’-*DDE, was performed by solid phase extraction using on-column degradation of the lipids and analysis by gas chromatography mass spectrometry at the Institut National de Santé Publique du Québec (INSPQ), Le Centre de Toxicologie du Québec, Canada (Table S1B). The detection limit was 200 ng/mL *p,p’-*DDE.

### DNA isolation for MethylC-Capture-seq (MCC-seq) on Greenlandic Inuit and South African sperm

Approximatively 10 million spermatozoa were lysed overnight at 37°C in a buffer containing a final concentration of 150 mM Tris, 10 mM ethylenediaminetetraacetic acid, 40 mM dithiothreitol, 2 mg/mL proteinase K, and 0.1% sarkosyl detergent. DNA was then extracted using the QIAamp DNA Mini kit (Qiagen) according to the manufacturer’s protocols. MCC-seq was performed as previously described (Chan et al., 2019), which involves an initial preparation of Whole Genome Bisulfite Sequencing (WGBS) libraries followed by targeted capturing of our regions of interest. Briefly, 1 – 2μg of DNA was sonicated (Covaris) and fragments of 300 – 400 bp were controlled on a Bioanalyzer DNA 1000 Chip (Agilent). Following this, the KAPA Biosystems’ protocols were used for DNA end repair, 3′-end adenylation, adaptor ligation, and clean-up steps. Using the Epitect Fast bisulfilte kit (Qiagen), samples were bisulfite converted according to manufacturer’s protocol, followed by quantification with OliGreen (Life Technology). PCR amplification with 9 – 12 cycles using the KAPA HiFi HotStart Uracil + DNA Polymerase (Roche/KAPA Biosystems) was performed according to suggested protocols. The final WGBS libraries were purified using Agencourt XP AMPure beads (Beckman Coulter), validated on Bioanalyzer High Sensitivity DNA Chips (Agilent), and quantified using PicoGreen (ThermoFisher). Following WGBS library preparations of all samples, regions of interest were captured using the SeqCap Epi Enrichment System protocol developed by RocheNimbleGen. Equal amounts of multiplexed libraries (84 ng of each; 12 samples per capture) were combined to obtain 1 μg of total input library and was hybridized at 47°C for 72h to the capture panel, specifically the human sperm capture panel recently developed by our group (Chan *et al*., 2019). This was followed by washing, recovery, PCR amplification of the captured libraries and final purification, according to manufacturer’s recommendations. Quality, concentration, and size distribution of the final captured libraries were determined using Bioanalyzer High Sensitivity DNA Chips (Agilent). The capture libraries were sequenced with a 200-cycle S2 kit (100 base paired-end sequencing) on the NovaSeq 6000 following the NovaSeq XP workflow (Table S2A,B).

### MCC-seq and data pre-processing

Targeted sperm panel MCC-Seq HiSeq reads were processed using the Genpipes pipeline (Bourgey et al., 2019). Specifically, the MCC-Seq paired-end fastq reads were first trimmed for quality (phred33 >= 30), length (n > 50bp), and removal of Illumina adapters using Trimmomatic (version 0.36) (Bolger et al., 2014). The trimmed reads were then aligned to the bisulfite-converted hg19 / GRCh37 reference genome with Bismark (version 0.18.1) (Krueger and Andrews, 2011) and Bowtie 2 (version 318 2.3.1) (Langmead and Salzberg, 2012) in paired-end mode using the non-directional protocol setting and the other default parameters. Bam files were merged and then de-duplicated with Picard (version 2.9.0). Methylation calls were obtained using Bismark to record counts of methylated and unmethylated cytosines at each cytosine position in the human genome. A methylation level of each CpG was calculated by the number of methylated reads over the total number of sequenced reads. CpGs that overlapped with SNPs (dbSNP 137) and the DAC Blacklisted Regions or Duke Excluded Regions (generated by the Encyclopedia of DNA elements - ENCODE project) (Amemiya et al., 2019) were removed. CpG sites with less than 20X read coverage were also discarded. Sperm sample purity (absence of somatic cell contamination) was assessed by examination of sequencing data for imprinted loci.

### Single nucleotide polymorphism and genotype analysis

BiSNP (version 0.82.2) (Liu et al., 2012) was run on the deduplicated bam files to call variants (including homozygous alternate and heterozygous genotypes). A set of unique variants called from all individuals were used as the potential SNP set. The homozygous reference genotypes of individuals on these SNPs were extracted from the aligned bam files by requiring >= 10X read coverage aligned to the reference allele. SNPs with genotypes inferred from all individuals were kept for downstream analysis. Principal component analysis on genotype profiles were used to investigate the genetic background distribution of samples used in this study. Chromosome 1 genotype data from 1,000 genomes (Imputed hapmap V3) were used as the reference genotype profile (the human population genetic background) to compare with that of South-African and Greenlandic cohorts.

### Differentially methylated CpGs (DMCs) and regions (DMRs) identification

Generalized linear regression models (GLMs) were built using the methylation proportion inferred from the combination of methylated reads and unmethylated reads as a binomially distributed response variable to look for associations between DNAme in sperm and *p.p’-*DDE exposure. Continuous and binarized *p.p’-*DDE effects were both explored and both models were adjusted for BMI, smoking status and age. For some CpGs, the number of individuals with sufficient sequencing coverage (>= 20X) was low (e.g. < 30 samples); these CpGs were removed from our analyses, to minimize the impact of low measurement accuracy. Non-variable CpGs (standard deviation = 0) were also removed to reduce the multiple testing burden. The R function glm() and the binomial family were used to fit each model, and p-values for variables of interest were obtained accordingly. The obtained p-values were then corrected by estimating the false discovery rate q-values using the qvalue R package. We defined significant associated DMCs when q-values were less than 0.01. Furthermore, consecutive DMCs that spanned 500 bp apart with methylation changes in the same direction were merged to define differentially methylated regions.

### MCC-seq hotspot cluster analysis

“Hotspot” or cluster analysis was performed by calculating the ratio of DMCs with DNAme gain or loss over the total number of CpGs found within 1 Mb sliding windows over the genome; densities >10% (termed clusters) were extracted for further analysis. To investigate genetic effects on DNAme of CpGs within the clusters, methylation quantitative trait locus (meQTLs) analyses were performed. The genotype profiles of SNPs within all the candidate clusters as well as all the DMCs were extracted. By considering possible SNP cis-effects within 250 kb of a CpG (i.e., a 500-kb window), meQTLs were calculated using MatrixEQTL (R package) with default parameters (Shabalin, 2012). The reported p-values were corrected using Benjamini-Hochberg false discovery rate (FDR) (Benjamini and Hochberg, 1995).

### Chromatin immunoprecipitation followed by sequencing (ChIP-seq) on South African sperm

Tubes containing South African sperm were thawed at room temperature and pelleted to remove freezing medium supernatant. Pellet was dissolved in 1 mL Ferticult Flushing medium. Samples were centrifuged, supernatant was removed, and an additional 500 µL of Ferticult Flushing medium was added to the sperm pellet. The sperm were incubated at room temperature for 45 minutes to allow for a swim-up. Fractions containing motile sperm were retrieved for downstream ChIP. ChIP on human sperm was performed as previously described (Hisano et al., 2013; Lambrot *et al*., 2021) with slight modifications. Sperm sample purity (absence of somatic cell contamination) was assessed visually in a hemocytometer counting chamber. Briefly, per sample, 12 million spermatozoa were resuspended in 300 mL of buffer 1 (15 mM Tris-HCl, 60 mM KCl, 5 mM MgCl2 and 0.1 mM EGTA) containing 0.3 M sucrose and 10 mM DTT. The samples were then each split into 6 tubes (2 million spermatozoa cells / tube) and 50 mL of buffer 1 was supplemented with 0.5% NP-40 and 1% sodium deoxycholate and added to each tube. After a 30-minute incubation on ice, 100 mL of MNase buffer (85 mM Tris-HCl, pH 7.5, 3 mM MgCl2 and 2 mM CaCl2) containing 0.3 M of sucrose and MNase (30 units of MNase for every 2 million sperm, Roche Nuclease S7) was added to each tube. The tubes were immediately placed at 37°C in a thermomixer for exactly 5 minutes. The MNase treatment was stopped by adding 2 mL of 0.5 M EDTA and placing the tubes on ice for 10 to 20 minutes. The tubes were then centrifuged at 17,000 x g for 10 minutes at room temperature to separate the debris and protamines (pellet) from the sheared chromatin (supernatant). Supernatants from the same sample were pooled into a 1.5 mL tube and 1X protease inhibitor (Roche) was added to the chromatin. The immunoprecipitation for H3K4me3 (Cell Signaling Technology) was carried out overnight at 4°C using Protein A Dynabeads (ThermoFisher Scientific). The mononucleosomal fraction (147 bp) was size selected using Agencourt XP AMPure beads (Beckman Coulter). Libraries were prepared using the Qiagen Ultralow Input Library Kit (Qiagen, #180495) as per manufacturer’s recommendations and samples were sequenced using paired-end 100 base pair reads with the NovaSeq 6000 platform (n = 49, Table S5A).

### ChIP-seq data pre-processing

Raw reads were trimmed with the TrimGalore wrapper script around the sequence-grooming tool cutadapt with the following quality trimming and filtering parameters (--length 50 -q 5 --stringency 1 -e 0.1’) (Krueger, 2015). The trimmed reads were mapped onto the hg19/GRCh37 reference genome downloaded from UCSC genome browser using bowtie2 as previously described (Lambrot *et al*., 2021; Langmead and Salzberg, 2012). We excluded reads that exhibited more than 3 mismatches. SAMtools was then used to convert SAM files and index BAM files. BigWig coverage tracks and binding heatmaps were generated from the aligned reads using deepTools2. The coverage was calculated as the number of reads extended to 150bp fragment size per 25 bp bin and normalized using Reads Per Kilobase per Million mapped reads (RPKM) not located on the X chromosome.

### H3K4me3 differential enrichment analysis

We previously identified H3K4me3 enrichment location in sperm using a high-quality reference human dataset (Lambrot *et al*., 2021). The exceptionally high sequencing depth of this dataset allowed us to obtain a robust and high-resolution map of H3K4me3 peaks in sperm (n = 50,117 peaks). We computed the sum of read counts under peaks, confirmed the robustness of the signals in the present dataset and excluded peaks with low counts (< 5 reads) in this study (n = 48,499 H3K4me3 peaks; Fig. S4A,B). Differential binding analysis was conducted to identify regions exhibiting different levels of H3k4me3 binding in sperm of individuals with the highest level of DDT exposure (third tertile, ter3; DDT > 14,102 ng / mL) compared to individuals with the lowest level exposure (first tertile, ter1; DDT < 1,132 ng / mL). To do so we used the edgeR Bioconductor/R package using the negative binomial distribution and shrinkage estimates of the dispersions to model read counts. Regions with FDR (Benjamini and Hochberg, 1995) below 0.2 were defined as significantly differentially enriched regions (deH3K4me3) between the two exposure groups.

### Dose-response analyses

For MCC-seq datasets, dose-response relationships were assessed via linear regressions conducted between the average percent DNAme at DNAme gain or loss DMCs relative to log10 serum *p,p’-*DDE concentration in Greenlandic and South African populations, respectively (Fig. 1C). For ChIP-seq datasets, dose-response analyses were assessed via Kolmogorov-Smirnov tests to compare the empirical cumulative distribution functions for regions with H3K4me3 gain or loss in sperm of South African men exposed to increasing categorical levels of *p,p’-*DDE, namely ter1, ter2 and ter3 (Fig. 3F,G).

**Fig. 1:**
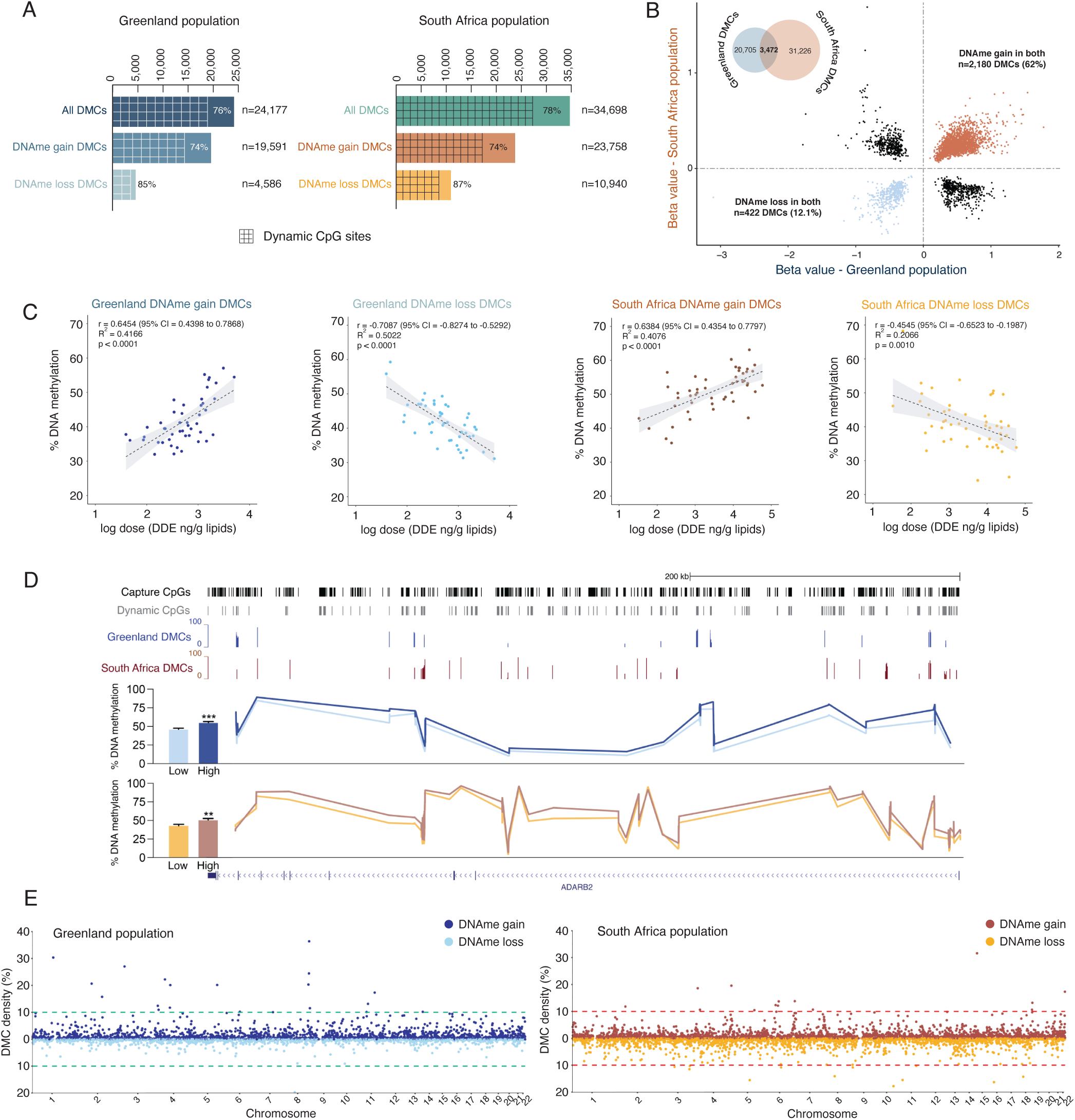
Exposure to *p,p’-*DDE appears to impact human sperm DNAme in a dose-dependent manner. (A) Number of differentially methylated sites (DMCs), DNAme gain DMCs, and DNAme loss DMCs, in the sperm of DDT-exposed Greenland or South African men. Number of dynamic DMCs (DNAme between 20 - 80%) are indicated by grids and percentages on the bar graphs. (B) Scatterplot of overlapping DMCs in Greenland and South African sperm (= 3,472 overlapping DMCs). Orange dots correspond to DMCs that gain DNAme in both Greenland and South African sperm (= 2,180 DMCs; 62% of total DMCs). Blue dots correspond to DMCs that lose DNAme in both Greenland and South African sperm (= 422 DMCs; 12.1% of total DMCs). (C) Average percent DNAme at DNAme gain or loss DMCs in Greenland or South African sperm relative to log10 serum *p,p’-*DDE concentration (in ng/g) for each individual. Linear regression line is plotted in dashed black and confidence interval in light grey. (D) Tracks at the ADARB2 locus showing percent DNAme levels in Greenland and South African sperm categorized based on low or high serum *p,p’*-DDE exposure levels (Greenland: low in light blue < 350 ng/g, n = 17 and high in dark blue > 900 ng/g, n = 18; South Africa: low in yellow < 1,200 ng/g, n = 19 and high in brown > 14,000 ng/g, n = 17). All CpGs captured by the MCC-seq are represented in black and dynamic CpGs in grey. CpGs captured in the Greenland sperm dataset are in blue and CpGs captured in the South African sperm dataset are red. (E) Manhattan plots on hotspot analysis for Greenland (blue) or South African (orange) sperm. Cluster analysis was performed by calculating the ratio of DMCs with DNAme gain or loss over the total number of CpGs found within 1 Mb sliding windows over the genome; densities > 10% (termed clusters) were extracted for further analysis.

### Transposable element and putative fetal enhancer annotations

For the transposable element analysis, hg19 RepeatMasker library was downloaded from http://www.repeatmasker.org/species/hg.html (hg19 - Feb 2009 - RepeatMasker open-4.0.5 - Repeat Library 20140131) and genomic ranges for classes of transposable and other repetitive DNA elements were catalogued. Genomic ranges for putative tissue specific fetal enhancers were retrieved from the Enhancer Atlas 2.0 (http://www.enhanceratlas.org) (Gao and Qian, 2020).

### Classification of transposable elements by % divergence score to infer transposable element age

hg19 RepeatMasker library includes divergence scores for each transposable element. For DNAme analysis, LTR- ERV1 (enriched at DMRs gaining and losing DNAme in Greenland and South African sperm) transposable elements described in the RepeatMasker library were sorted into percent divergence quantiles (Fig. S3G). For H3K4me3 analysis, LINE-1, LTR-ERV1-MaLR (enriched at H3K4me3 gaining regions in South African sperm), LINE-2, SINE-MIR, and SINE-Alu (enriched at H3K4me3 losing regions in South African sperm) transposable elements described in the RepeatMasker library were similarly sorted into percent divergence quantiles (Fig. S4G – K). Transposable elements belonging to the 1st quarter were characterized as young, 2nd quarter as mid-young, 3rd quarter as mid-old, and 4th quarter as old (Lanciano and Cristofari, 2020).

### Enrichment and gene ontology analyses on DMRs and deH3K4me3 peaks

The significance of functional gene ontology (GO) enrichment was estimated using a weighted fisher test (“weight01” algorithm) as implemented in the R/Bioconductor topGO package (Alexa et al., 2006). Significant overlaps of DMRs or deH3K4me3 with specific genomic location (including genic annotation, overlapping specific families of transposable elements or putative enhancers) were calculated using a permutation test framework implemented in the R/Bioconductor regioneR package (Gel et al., 2016). Random regions (of the same size than the tested regions) were resampled from the background list of MCC-seq regions or H3K4me3 peaks. In Fig. 5D – F, we asked whether regions that bear both changes in DNAme and H3K4me3 overlapped specific genomic location compared to regions that bear changes in only DNAme or H3K4me3. In that case, random regions were resampled from the list of DMR or deH3K4me3, respectively.

### Identification of DNAme sperm-to-embryo persistent regions

RRBS pre-implantation embryo datasets were retrieved from Guo et al., 2014 (GSE49828). Sperm DNAme CpGs were classified into low (< 20% DNAme), dynamic (between 20 - 80% DNAme), or high (> 80% DNAme) DNAme levels and compared to pre-implantation embryo CpG DNAme levels. DNAme sperm-to-zygote persistent CpGs retained the same levels of DNAme from sperm to the zygote whereas DNAme sperm-to-ICM persistent CpGs retained the same levels of DNAme from sperm to the ICM. Low, dynamic, or high DNAme persistent CpGs that spanned 500 bp apart were then merged to generate DNAme sperm-to-zygote and sperm-to-ICM persistent regions. We identified 18,562 low, 2,132 dynamic and 5,745 high DNAme sperm-to-zygote persistent regions, and 14,235 low, 488 dynamic and 694 high DNAme sperm-to-ICM persistent regions. Significant overlaps od DNAme sperm-to-embryo persistent regions with specific genomic location (including genic annotation, overlapping specific families of transposable elements or putative enhancers) were calculated using a permutation test framework implemented in the R/Bioconductor regioneR package. Random regions (of the same size than the tested regions) were resampled from the background list of MCC-seq regions overlapping RRBS regions profiled in Guo et al., 2014.

### Pre-implantation embryo H3K4me3 dataset pre-processing

H3K4me3 ChIP-seq pre-implantation embryo raw reads were retrieved from Xia et al., 2019 (GSE124718). Reads were trimmed using Trimmomatic in paired-end mode (version 0.38) (Bolger et al., 2014). The trimmed reads were mapped onto the hg19/GRCh37 reference genome downloaded from UCSC genome browser using bowtie2 as previously described (Lambrot *et al*., 2021).

### Association between promoter sperm, embryo H3K4me3 counts, and embryo gene expression

To identify promoters that were enriched for sperm, 4-cell, 8-cell, and ICM H3K4me3, we plotted the density distribution of the log2 (mean counts + 8) and identified the cutoff value between lowly and highly abundant signal based on the local minimum of the bimodal distribution (Fig. S6D – G). Of note, 4-cell embryo H3K4me3 density distribution did not follow a bimodal trend therefore the cutoff value between low and high abundant signal was determined by visually assessing the distribution of counts against its density (Fig. S6E). In each embryo stage, genes with an RPKM > 1 were considered expressed.

### Pre-implantation embryo H3K4me3 enrichment

Regions enriched for H3K4me3 in 4-cell embryos, 8-cell embryos, and ICM, were identified using the R/Bioconductor package csaw (Lun and Smyth, 2016). Reads with mapping quality score above 20 were counted in 150 bp sliding windows for each library across the genome after exclusion of blacklisted regions (Amemiya *et al*., 2019). To estimate global background signal, reads were counted in 2,000 bp contiguous bins for each library across the genome. We then identified regions enriched for H3K4me3 by filtering windows with background (non-specific) enrichment, and by merging contiguous 150 bp windows that were remaining. All parameters were optimized independently for each stage of pre-implantation embryogenesis after visual assessment of tracks using Integrative Genome Viewer (IGV). Windows with a log2 fold change over 7 (for H3K4me3 4-cell embryo data), over 18 (for H3K4me3 8-cell embryo data), or over 6 (for H3K4me3 ICM data) were merged. Maximum peak size was set at 20,000 bp for all embryo stages. This yielded 42,630, 25,457, and 18,284 H3K4me3 peaks in 4-cell embryos, 8-cell embryos, and ICM respectively.

### Identification of H3K4me3 sperm-to-embryo persistent peaks

To identify persistent sperm-to-4-cell H3K4me3 peaks, we overlapped sperm H3K4me3 peaks to 4-cell H3K4me3 peaks. To identify persistent sperm-to-ICM H3K4me3 peaks, we first overlapped persistent sperm-to-4-cell H3K4me3 peaks to 8-cell H3K4me3 peaks to generate persistent sperm-to-8-cell H3K4me3 peaks. Then, we overlapped persistent sperm-to-8-cell H3K4me3 peaks to ICM H3K4me3 peaks which yielded sperm-to-ICM H3K4me3 peaks.

## Results

### *p,p’-*DDE exposure and genetic diversity in Greenlandic Inuit and South African VhaVenda Men

Sperm samples from men of two geographically distinct populations exposed to DDT or its metabolite *p,p’-*DDE were used for this study to investigate the association between exposure and DNAme and / or histone methylation in sperm (see Methods for participant details). To study the consequences of *p,p’-*DDE bioaccumulation in Northern populations on sperm DNAme and H3K4me3, we used 47 paired serum and semen samples from Greenlandic Inuit men of the INUENDO cohort (Fig. S1A and Table S1A) (Bonde et al., 2008; Spanò *et al*., 2005; Toft et al., 2005). The men ranged from 20 to 44 years of age with a mean age of 31 years, and had serum levels of the DDT metabolite, *p,p’-*DDE, between 39.4 to 5,000 ng/g lipid (Fig. S1A and Table S1A).

To investigate the relationship between direct exposure to DDT through indoor spraying and the sperm epigenome, we used 49 semen samples from a cohort of men who were recruited from 12 villages in the Thulamela Local Municipality, in the Vhembe District Municipality of Limpopo Province, that were either sprayed or not (Fig. S1B, Table S1B). The men were between 18 to 32 years of age with a mean age of 25 years. Body burdens of *p,p’-*DDE ranged from 33 to 58,544 ng/g of lipids with a mean of 32,000 ng/g of lipids (Fig. S1B and Table S1B). Sample selection was based on sperm quality requirements for ChIP-seq (normal DNA fragmentation index and sperm count > 10 million). In comparison to the Greenlandic Inuit men, the South African VhaVenda men had on average 12 times higher serum *p,p’-*DDE levels (Table S1).

To exclude genetic variation as a potential source of epigenetic variation, we assessed the genetic diversity of the Greenlandic and South African populations (see Methods). Principal component analysis (PCA) on genotype profiles of Greenlandic Inuit and South African VhaVenda cohorts compared to human reference populations, showed unique population ancestries for both cohorts while demonstrating the population homogeneity within each cohort (Fig. S1C). Consequently, the results indicate that the likelihood of ethnic subgroups being the source of observed epigenetic variation is low.

### Serum *p,p’-*DDE levels in Greenlandic and South African men are associated with dose-response alterations in sperm CpG methylation

Sperm DNA methylation was assessed using MCC-seq, targeting a select set of regions and providing sequencing-based information on millions of CpG sites. Specifically, we used the human 5-methylcytosine-capture sequencing sperm capture panel recently developed by our group (Chan *et al*., 2019). This panel interrogates approximatively 3.18 million CpGs in the genome, including the > 850,000 sites present on the Infinium MethylationEPIC BeadChip, an array-based technique targeting commonly assessed gene promoter / CpG island regions and enhancers, and widely used in DNA methylation analyses. In addition, the human sperm capture panel targets approximately one million dynamic CpGs, that are environmentally sensitive sequences which demonstrate higher variability and possess intermediate levels of methylation (between 20 – 80%) in human sperm. The quantitative association between sperm DNAme and serum *p,p’-*DDE was assessed using a generalized linear regression model adjusted for potential confounders (smoking, age, and body mass index; Table S1A,B and S2A,B). We applied a continuous analysis to identify differentially methylated cytosines (DMCs) with DNAme gain or loss, and identified 24,177 differentially methylated CpGs in Greenlandic sperm and 34,698 DMCs in South African sperm (Fig. 1A; Table S2C,D). In both populations, over 75% of DMCs were found at dynamic CpGs (methylation levels between 20 and 80%; Fig. 1A). DMCs with a DNAme gain were more abundant than DMCs with a DNAme loss in Greenlandic and South African sperm (19,591 and 23,758 DMCs with a DNAme gain and 4,586 and 10,940 DMCs with a DNAme loss respectively; Fisher test p < 0.0001; Fig. 1A). Interestingly, 3,472 DMCs overlapped between both populations (Fisher test p < 0.0001) with 62% of the overlapping DMCs gaining DNAme (= 2,180 DMCs; Fisher test p < 0.0001; Fig. 1B). Previous studies have shown that DNAme across various cell types exhibit a dose-response to toxicants such as arsenic and cigarettes (Niedzwiecki Megan et al., 2013; Zhang et al., 2016). To determine whether DNAme in sperm was dose-responsive to *p,p’-*DDE, we investigated the linear relationship between *p,p’-*DDE from men’s serum and average changes in DMCs from their paired sperm samples (Fig. 1C). Percent DNAme level at DMCs in Greenlandic or South African sperm, and serum *p,p’-*DDE values, showed a linear dose-response trend (Fig. 1C; r = 0.6454 for Greenland DNAme gain DMCs, -0.7087 for Greenland DNAme loss DMCs, 0.6384 for South Africa DNAme gain DMCs, -0.4545 for South Africa DNAme loss DMCs, and p < 0.0001) across all categories.

An example of dose response difference in DNAme is shown for a gene of interest, adenosine deaminase, RNA-Specific, B2 (ADARB2) (Fig. 1D). ADARB2 is important for the regulation of RNA editing (Nishikura, 2010) and its perturbed function has been implicated in childhood cancer and neurological diseases (Akbarian et al., 1995; Paz et al., 2007). A total of 162 and 160 DMCs were identified throughout the ADARB2 locus in the Greenlandic and South African sperm respectively, of which 32 DMCs were common between both populations (Fig. 1D). The DMCs at the ADARB2 locus were predominantly at dynamic CpGs and displayed intermediate levels of DNAme (129 and 132 dynamic DMCs in the Greenlandic and South African samples respectively) (Fig. 1D). We categorized samples based on low or high *p,p’-*DDE exposure levels (Greenland: low < 350 ng/g, n = 17 and high > 900 ng/g, n = 18; South Africa: low < 1,200 ng/g, n = 19 and high > 14,000 ng/g, n = 17). Subjects with high *p,p’-*DDE exposure had higher levels of sperm DNAme than those with low exposure across the ADARB2 locus (t-test, p < 0.05; Fig. 1D).

As we previously reported for age-related alterations in sperm DNAme, DMCs can be found densely clustered in some areas of the genome, suggesting the possibility of genomic “hotspots” for environmental exposures (Cao et al., 2020). We identified several chromosomal regions of high DMC density (over 10%; termed clusters), in the sperm of both populations that were not confounded by genetic variations (Fig. 1E; Table S2E,F). A total of 21 clusters of differential methylation were identified in the Greenlandic sperm (20 with DNAme gain, 1 with DNAme loss; Table S2E), and 26 in the South African sperm (14 with DNAme gain, 12 with DNAme loss; Table S2F). Two DNAme gain and 1 DNAme loss clusters were common between both cohorts. In the South African population, a cluster containing 443 DMCs overlapped to the imprinting control SNRPN region, which is implicated Prader Willi and Angelman syndromes (Reed and Leff, 1994) (Table S2F). Many DMCs were also annotated to the SNORD family of genes, which have been shown to have altered DNAme in sperm samples from the Early Autism Risk Longitudinal Investigation cohort (Feinberg et al., 2015) (Table S2E,F).

### Differentially methylated regions in Greenlandic and South African sperm are implicated in development and nervous system function

To gain functional insight into how *p,p’-*DDE associated alterations in sperm DNAme may impact sperm function and embryo development, differentially methylated regions (DMRs) were identified and their relationship to genome function and embryo development was explored. A DMR was defined as a region with two or more DMCs with a consistent DNAme gain or loss and spanning a maximum of 500 bp apart (DMRs; Fig. 2A; Table S2G,H). This merging of DMCs yielded 6,787 DMRs in Greenlandic sperm and 12,849 DMRs in South African sperm (Fig. 2A; Table S2G,H). Overall, 1,187 DMRs in Greenland sperm overlapped DMRs in South African sperm and 1,189 DMRs in South African sperm overlapped DMRs in Greenlandic sperm (Fig. 2B). In line with the DMC analysis, DMRs with increased DNAme were more abundant than DMRs with decreased DNAme in both populations (4,928 DNAme gain DMR in Greenlandic sperm and 8,259 DNAme gain DMR in South African sperm; Fig. 2C). DMRs were predominantly enriched at > 10 kb from a transcriptional start site (TSS) in intergenic space for both populations (Fisher test p < 0.0001; Fig. 2D and Fig. S2A). Gene ontology analysis on genes that overlapped a sperm DMR revealed common significant pathways in both populations, highlighting the consistency of *p,p’-*DDE-associated effects on the sperm epigenome across different populations. Common significant pathways were involved in the development and function of the nervous system such as axon guidance, regulation of synapse assembly and neurogenesis (Fig. 2E – H; Table S3A – D). These pathways included genes *DVL1*, *ROBO1* and *TRIO.* Notably, all are implicated in neurodevelopmental disorders such as autism and dyslexia which have been reported as adverse outcomes from developmental exposures to endocrine disrupting chemicals (EDC) (Cheroni et al., 2020; World Health et al., 2013). Developmental processes were also identified and impacted common genes in both populations such as *PARD3* (neural tube defects), *MYLK* (aorta development) and *HOXB2* (skeletal patterning) (Fig. 2E – H; Table S3A – D).

**Fig. 2:**
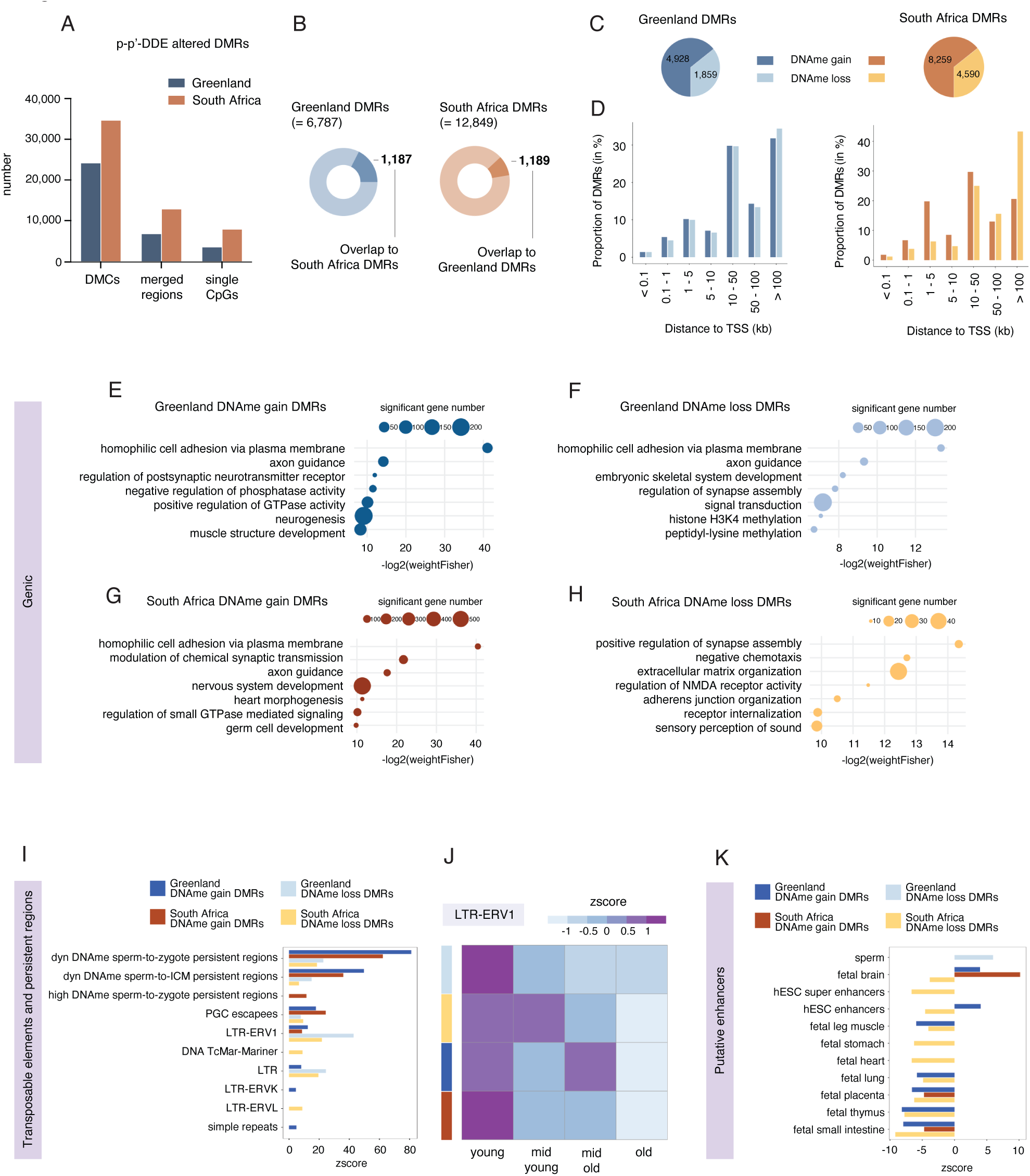
Differentially methylated regions are involved in development and intersect young retrotransposons that retain DNAme during embryogenesis. (A) Number of DMCs, differentially methylated regions (DMRs), and single CpGs in Greenland (blue) and South African (orange) sperm. DMRs were called by merging DMCs separated by a maximum distance of 500 bp. (B) Proportion of Greenland sperm DMRs (= 6,787) that overlap South African sperm DMRs (overlap = 1,187) and South African sperm DMRs (= 12,849) that overlap Greenland sperm DMRs (overlap = 1,189). (C) Number of DNAme gain or loss DMRs in Greenland or South African sperm. (D) Genomic distribution of DMRs with DNAme gain or loss in Greenland or South African sperm relative to the transcriptional start site (TSS). (E - H) Selected significant pathways from gene ontology analysis on genes at a DMR with DNAme gain or loss in Greenland or South African sperm (weighed Fisher p < 0.05). Size of circle corresponds to the number of genes from a significant pathway that overlap a DMR. (I) Enrichment of DNAme gain or loss DMRs in Greenland or South African sperm at identified DNAme embryonic persistent regions (see Fig. S3, primordial germ cell (PGC) escapees, and transposable element annotations (RepeatMasker hg19 library 20140131). Positive enrichments are determined by Z scores using the Bioconductor package regioneR. For all annotations displayed, p < 0.0001 and n = 10,000 permutations of random regions (of the same size) resampled from the targeted MCC-seq regions. (J) Distribution of DNAme gain or loss DMRs in Greenland or South African sperm relative to age quarters of LTR- ERV1 transposable elements (significantly enriched in the four DMR categories; Fig. 2H). Age of LTR-ERV1 transposable elements was determined by partitioning the transposable elements’ percent divergence scores into quarters where first quarter = low percent divergence and young LTR-ERV1; second quarter = mid-low percent divergence and mid-young LTR-ERV1; third quarter = mid-high percent divergence and mid-old LTR-ERV1; fourth quarter = high percent divergence and old LTR-ERV1 (see Fig. S3B). (K) Enrichment of DNAme gain or loss DMRs in Greenland or South African sperm at putative tissue-specific enhancer classes. For all annotations displayed, p < 0.0001 and n = 10,000 permutations of random regions (of the same size) resampled from the targeted MCC-seq regions.

### Predicted sperm-to-embryo persistent DNAme regions include DMRs associated with *p,p’-*DDE exposure

We then aimed to investigate whether exposure to *p,p’-*DDE was associated with sperm DMRs that are predicted to persist in the embryo and could therefore potentially be implicated in epigenetic inheritance. To do so, we performed a stepwise analysis initially focused on identifying sperm transmitted DNAme that potentially escapes reprogramming. First, we identified CpGs that retained the same level of DNAme from sperm to the zygote and from sperm to the inner cell mass (ICM) of the blastocyst using our data and an existing data set (Guo *et al*., 2014). CpGs that retained sperm- specific DNAme patterns in the embryo may correspond to CpGs that escape reprogramming (Fig. S3 and Table S4A – F). By this approach we identified: 1) CpGs with DNAme that is consistent from sperm to the zygote corresponding to CpGs that could influence the first wave of zygotic gene expression (Fig. S3A – B); and 2) CpGs with DNAme that persist across pre-implantation embryogenesis until the first lineage specification in the blastocyst (Fig. S3C – D).

We then merged two or more low, dynamic, or high DNAme persistent CpGs spanning a maximum of 500 bp apart to classify DNAme sperm-to-zygote and sperm-to-ICM persistent regions (Fig. S3A – D; Table S4A – F). This yielded 18,562 low, 2,132 dynamic and 5,745 high DNAme sperm-to-zygote persistent regions (Fig. S3B; Table S4A – C), and 14,235 low, 488 dynamic and 694 high DNAme sperm-to-ICM persistent regions (Fig. S3D; Table S4D – F). Dynamic and high DNAme persistent regions were enriched at CpG shores, exons, CpG shelves, and introns (Fig. S3E; z-score > 0; p < 0.0001; n = 10,000 permutations). Dynamic DNAme persistent regions were more enriched in intergenic space than high DNAme persistent regions (Fig. S3E). Low DNAme persistent regions were not enriched at any genic or CpG annotations (Fig. S3E). Remarkably, a significant proportion of sperm-to-zygote and sperm-to-ICM persistent regions annotated to transposable elements (TEs) with the LTR-ERV1, LINE-2, and LTR-ERVL-MaLR TE families being over-represented across dynamic and high DNAme persistent regions (Fig. S3F; z-score > 0; p < 0.0001; n = 10,000 permutations). Analysis of functional elements also revealed an enrichment of human embryonic stem cell (ESC) enhancers and super enhancers at dynamic and high DNAme persistent regions (Fig. S3F; z-score > 0; p < 0.0001; n = 10,000 permutations) (Barakat et al., 2018).

Lastly, to identify *p,p’-*DDE-associated DMRs that may be epigenetically inherited, we determined if DMRs co-localized to DNAme persistent regions (Fig. 2I). Dynamic DNAme sperm-to-zygote and sperm-to-ICM persistent regions, as well as previously described PGC escapees (Tang et al., 2015), were enriched across all DMR categories (Fig. 2I; z-score > 0; p < 0.0001; n = 10,000 permutations), supporting the possibility that dynamic DNAme persistent regions impacted by *p,p’-*DDE may escape reprogramming in the embryo. DMRs were also enriched for certain TE families, with an overrepresentation of the LTR-ERV1 family at DMRs with DNAme gain and loss in Greenlandic and South African sperm (Fig. 2I). The age of a TE can be inferred from the level of divergence observed between the TE sequence and the canonical full-length element sequence (Schmidt et al., 2012). Indeed, this value reflects the duration the TE has been integrated into the genome, and thus an indicator of TE age (Schmidt et al., 2012). Importantly, more newly integrated young TEs are typically more active than old TEs (Lanciano and Cristofari, 2020). Consequently, epigenetic changes at more recently integrated TEs may elicit more dramatic effects on gene expression. We categorized all LTR-ERV1 TEs of the human genome into four quarters based on their percent divergence scores (Fig. S3G), Interestingly, young LTR-ERV1s were the predominant age quarter that overlapped DMRs (Fig. 2J and Fig. S3H). Because the majority of DMRs were found in intergenic space, we intersected them to putative fetal tissue enhancers (Fig. 2K). DMRs with DNAme gain in Greenland and South African sperm were overrepresented at fetal brain putative enhancers whereas DMRs with DNAme loss in Greenland sperm were enriched at sperm putative enhancers (Fig. 2K; z-score > 0; p < 0.0001; n = 10,000 permutations) (Gao and Qian, 2020). All other studied classes of putative fetal enhancers were underrepresented at DMRs, highlighting the functional specificity of *p,p’*-DDE-associated regions in fertility and neurodevelopment (Fig. 2K). Taken together the analysis of MCC-Seq data suggests that dynamic DNAme CpGs in sperm are more sensitive to alterations associated with *p,p’*- DDE exposure and may escape epigenetic reprogramming in the embryo at young transposable elements.

### *p,p’-*DDE exposure is associated with altered sperm H3K4me3 enrichment at important developmental regulatory loci

Next, we set out to investigate whether exposure to *p,p’-*DDE was associated with differential enrichment of H3K4me3 (deH3K4me3) in human sperm and if changes occurred at genomic regions associated with the observed population health abnormalities in DDT-exposed regions. To do so, we performed 100 bp paired-end sequencing on high-quality H3K4me3 chromatin immunoprecipitation libraries from the sperm of each participant of the South African VhaVenda cohort. On average, 98% of reads aligned to the human genome, yielding approximately 115 million reads per sperm sample, of which 88 million were uniquely mapped (Table S5A). We identified 48,499 H3K4me3 peaks in South African sperm samples (Fig. S4A and B). Sperm samples were categorized based on tertiles of their corresponding serum *p,p’*-DDE levels as low (Ter1), intermediate (Ter2) or high (Ter3) exposure (Fig. S1B), and a differential enrichment analysis was performed to identify differential H3K4me3 peaks between categories of *p,p’*-DDE levels (Lun and Smyth, 2016). Comparison between high (Ter3) versus low (Ter1) *p,p’*-DDE levels identified 1,865 peaks with differentially enriched H3K4me3 (deH3K4me3; FDR < 0.2) of which 851 peaks gained H3K4me3 enrichment and 1,014 peaks lost H3K4me3 enrichment (Fig. 3A; Table S5B). We did not observe significant differences after adjustment for BMI or age and therefore did not include these variables in our models. Peaks gaining H3K4me3 were enriched in intergenic regions (Fig. S4C; z-score > 0; p < 0.0001; n = 10,000 permutations) and predominantly > 10 kb from the TSS (Fig. 3B). Conversely, peaks with H3K4me3 loss were primarily genic occurring at promoters, exons and at CpG rich regions (Fig. S4D; z-score > 0; p < 0.0001; n = 10,000 permutations), and mostly < 5 kb from the TSS (Fig. 3C). This suggests that peaks with H3K4me3 loss are more likely to directly impact gene expression in the next generation (Fig. 3C).

In rodents, H3K9ac was identified as dose-responsive to traffic air pollution in lung and blood cells (Ding et al., 2017). Whether sperm chromatin in humans shows a dose-response trend to any toxicant is unknown. To investigate this possibility, we first depicted the average H3K4me3 enrichment in deH3K4me3 peaks across the three levels of exposure. Consistent with a dose response effect, H3K4me3 enrichment were most apparent for the extreme levels of exposure at deH3K4me3 (Ter3 vs Ter1; Fig. 3D – G and Fig. S1B). Indeed, testing altered regions for a dose- like response across all exposure categories revealed significant effects for regions that gained H3K4me3 (Ter1 vs Ter2; Kolmogorov-Smirnov p < 0.0001 and Ter 2 vs Ter 3; Kolmogorov-Smirnov p < 0.05; Fig. 3F and Fig. S4E), and for regions that lost H3K4me3 (Ter1 vs Ter2 and Ter2 vs Ter3; Kolmogorov-Smirnov p < 0.0001; Fig. 3G and Fig. S4F). Examples of peaks with a dose-respond trend across the three tertiles were *FSIP1* gaining in H3K4me3 (important for acrosomal reaction and sperm flagellation; Fig. 3H) (Gamallat et al., 2021), and *BRD1* losing in H3K4me3 chromatin interacting protein involved in brain development; Fig. 3I) (Severinsen et al., 2006).

**Fig. 3:**
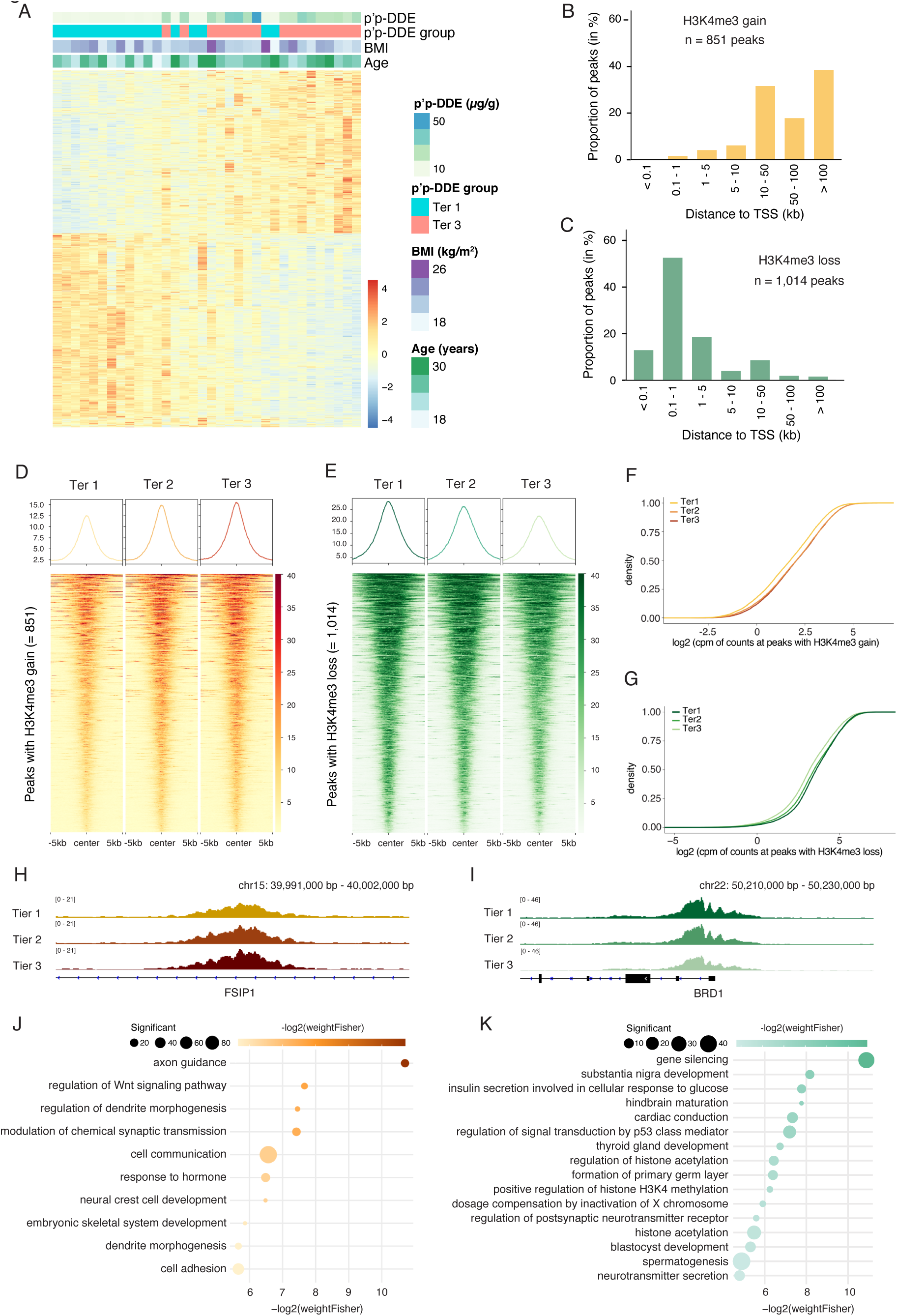
H3K4me3 is differentially enriched in sperm of South African men exposed to *p,p’-*DDE. (A) Heatmap of normalized H3K4me3 counts at the 1,865 peaks with differentially enriched H3K4me3 (deH3K4me3; 851 peaks with H3K4me3 gain and 1,014 peaks with H3K4me3 loss; FDR < 0.2) in sperm from South African men with low (tier 1) or high (tier 3) serum *p.p-*DDE levels. Serum *p.p-*DDE concentration, tier, body mass index (BMI), and age of participants is indicated by coloured boxes above the heatmap. (B) Genomic distribution of peaks with H3K4me3 gain in South African sperm relative to the TSS. (C) Genomic distribution of peaks with H3K4me3 loss in South African sperm relative to the TSS. (D - E) H3K4me3 signal intensity heatmaps at +/- 5 kb the center of the peaks with H3K4me3 gain in South African sperm (= 851 peaks) relative to the three *p.p-*DDE tier levels. (E) H3K4me3 signal intensity heatmaps at +/- 5 kb the center of the peaks with H3K4me3 loss in South African sperm (= 1,014 peaks) relative to the three *p.p-*DDE tier levels. (F) Estimator of the cumulative distribution function (ECDF) plot for log2 cpm of H3K4me3 counts at regions with H3K4me3 gain in tier1 (yellow), tier2 (orange), tier3 (red) sperm samples. Dose-response trends were assessed by via Kolmogorov-Smirnov tests with p < 0.0001 for ter1 vs ter2 at H3K4me3 gain regions, p < 0.05. (G) Estimator of the cumulative distribution function (ECDF) plot for log2 cpm of H3K4me3 counts at regions with H3K4me3 gain in tier1 (dark green), tier2 (medium green), tier3 (light green) sperm samples. Dose-response trends were assessed by via Kolmogorov-Smirnov tests with p < 0.0001 for ter1 vs ter2 at H3K4me3 gain regions, p < 0.0001. (H) Representative Integrative Genome Viewer (IGV) tracks of peak with H3K4me3 gain in South African sperm at the FSIP1 genic region. (I) Representative IGV tracks of peak with H3K4me3 loss in South African sperm at the BRD1 promoter. (J - K) Selected significant pathways from gene ontology analysis on genes with H3K4me3 gain (J) or loss (K) in South African sperm (weighed Fisher p < 0.05). Size of circle corresponds to the number of genes from a significant pathway that overlap a peak with H3K4me3 gain. Shade intensity of the circle indicates -log2(weightFisher) value of significant pathway.

### Peaks with gain or loss in H3K4me3 localize to specific families of transposable elements and gene regulatory regions

Gene ontology analyses on genes overlapping peaks with H3K4me3 gains revealed enrichment at genes involved in neural development, and cell signaling including hormone responses (Fig. 3J; Table S3E). Peaks with H3K4me3 loss were associated with genes and tissues implicated in metabolism, development, spermatogenesis, chromatin remodeling, and the endocrine system (Fig. 3K; Table S3F). Genes of interest for their roles in development and disease included: *NRP1*, *SOX6*, *SEMA5A* at peaks with H3K4me3 gain, and *KDM6B*, *SOX17*, *BRCA1* at peaks with H3K4me3 loss. Sperm-transmitted intergenerational effects of DDT exposure may be mediated by alterations of H3K4me3 at TEs in sperm. In fact, TEs that escape epigenetic reprogramming and that are regulated by chromatin features including H3K4me3, may behave as enhancers and / or promoters in the embryo (Chuong et al., 2013; Jacques et al., 2013; Lynch et al., 2011; Lynch et al., 2015; Veselovska et al., 2015). We identified a high degree of specificity in the TE families that overlapped peaks with gain or loss of H3K4me3 in sperm (Fig. 4A and B). Peaks with H3K4me3 gain were significantly enriched at LINE-1 elements, DNA repeats, and LTR-ERVL-MaLR (z-score > 0, p < 0.0001, n = 10,000 permutations; Fig. 4A). Conversely, TE and repeat families overrepresented at peaks with H3K4me3 loss included low complexity repeats, SINE-Alu (such as SINE-FLAM-A, SINE-FLAM-C, and SINE-FRAM), SINE-MIR, and LINE-2 (z-score > 0, p < 0.0001, n = 10,000 permutations; Fig. 4B).

**Fig. 4:**
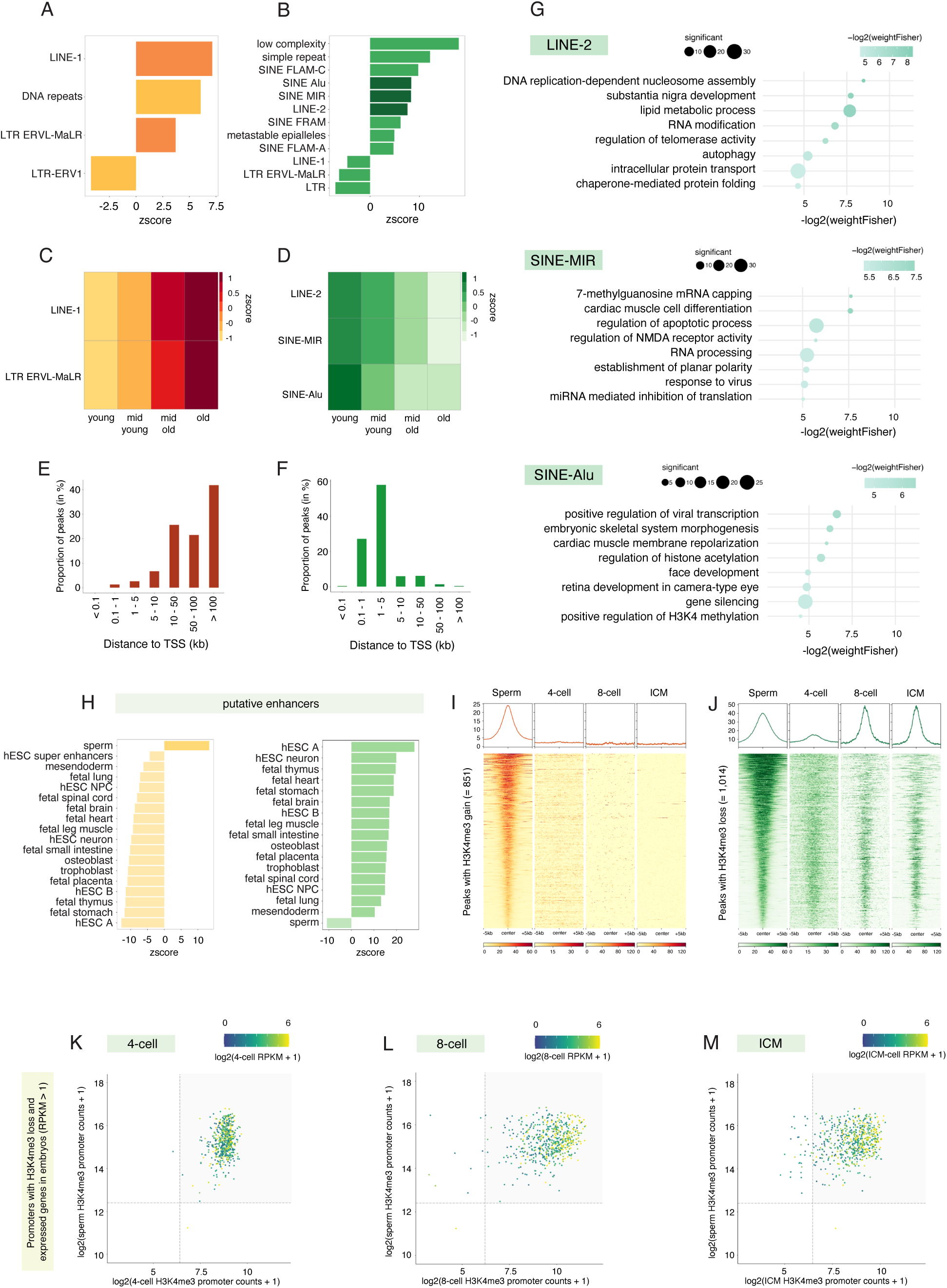
Peaks with H3K4me3 loss in South African sperm are enriched at young retrotransposons that retain H3K4me3 in the pre-implantation embryo. (A) Enrichment for peaks with H3K4me3 gain in South African sperm at transposable element annotations (RepeatMasker hg19 library 20140131). Positive and negative enrichments are determined by Z scores using the Bioconductor package regioneR. For all annotations displayed, p < 0.0001 and n = 10,000 permutations of random regions (of the same size) resampled from sperm H3K4me3 peaks. Enrichments coloured in dark orange correspond to functional transposable element classes used in Fig. 4C. (B) Enrichment for peaks with H3K4me3 loss in South African sperm at transposable element annotations (RepeatMasker hg19 library 20140131). Positive and negative enrichments are determined by Z scores using the Bioconductor package regioneR. For all annotations displayed, p < 0.0001 and n = 10,000 permutations of random regions (of the same size) resampled from sperm H3K4me3 peaks. Enrichments coloured in dark green correspond to functional transposable element classes used in Fig. 4D. (C) Distribution for peaks with H3K4me3 gain in South African sperm relative to age quarters of LINE-1 and LTR ERV-MaLR transposable element classes (significantly enriched for peaks with H3K4me3 gain; Fig. 4A). Age of transposable elements was determined by partitioning the class percent divergence score in quarters (see Fig. SG,H). (D) Distribution for peaks with H3K4me3 loss in South African sperm relative to age quarters of LINE-2, SINE-MIR, and SINE-Alu transposable element classes (significantly enriched for peaks with H3K4me3 loss; Fig. 4B). Age of transposable elements was determined by partitioning the class percent divergence score in quarters (see Fig. S4I- K). (E) Genomic distribution of peaks with H3K4me3 gain in South African sperm that overlap an old (4th quarter) LINE- 1 (= 139) or LTR ERVL-MaLR (= 74) transposable element, relative to the TSS. (F) Genomic distribution of peaks with H3K4me3 loss in South African sperm that overlap a young (1st quarter) LINE- 2 (= 187), SINE-MIR (= 267), or SINE-Alu (= 507) transposable element, relative to the TSS. (G) Selected significant pathways from gene ontology analysis on promoter peaks with H3K4me3 loss in sperm that overlap a young (1st quarter) LINE-2 (= 324 promoters), SINE-MIR (= 442 promoters), or SINE-Alu (= 809 promoters) transposable element at (weighed Fisher p < 0.05). Size of dots corresponds to the number of genes from a significant pathway that overlap a peak with H3K4me3 loss. Color of the dots indicates -log2(weightFisher) value of significant pathway. (H) Enrichment for peaks with H3K4me3 gain (yellow) or loss (green) in South African sperm at tissue-specific putative enhancer annotations. For all annotations displayed, p < 0.0001 and n = 10,000 permutations of random regions (of the same size) resampled from sperm H3K4me3 peaks. (I) Sperm and pre-implantation embryo (4-cell, 8-cell, ICM) H3K4me3 signal intensity heatmaps at +/- 5 kb the center of the peaks with H3K4me3 gain in South African sperm (= 851 peaks). (J) Sperm and pre-implantation embryo (4-cell, 8-cell, ICM) H3K4me3 signal intensity heatmaps at +/- 5 kb the center of the peaks with H3K4me3 loss in South African sperm (= 1,014 peaks). (K - M) Scatterplots where the x axis corresponds to the log2 (pre-implantation embryo H3K4me3 promoter counts + 1) and the y axis corresponds to the log2 (sperm H3K4me3 promoter counts + 1) at promoters with H3K4me3 loss in South African sperm that are expressed at the described stages of pre-implantation embryo development (RPKM > 1). Color of the scatter points correspond to the log2 pre-implantation embryo RPKM gene expression + 1. Dashed lines correspond to H3K4me3 promoter density cutoffs for pre-implantation embryo (x axis) or sperm (y axis). Grey box denotes promoters with H3K4me3 in sperm and 4-cell (K), 8-cell (L), and ICM (M) pre-implantation embryos.

Complementing the DNAme TE analysis, we selected the families of TEs that were over-represented in peaks with gain or loss of H3K4me3 and categorized them into young, mid-young, mid-old, and old based on their percent divergence scores (see Methods, Fig. S4G – K). Regions that gained H3K4me3 in sperm at LINE-1 and LTR ERVL- MaLR transposable elements, predominantly overlapped older TEs and were located > 10 kb from the TSS (Fig. 4C and E). In contrast, LINE-2, SINE-MIR and SINE-Alu that intersected peaks that loss H3K4me3 in sperm, were classified as younger TEs and mostly enriched at < 5 kb from the TSS (Fig. 4D and F). Next, we performed a gene ontology analysis for young LINE-2, SINE-MIR, and SINE-Alu elements that overlapped a gene with H3K4me3 loss in sperm (Fig. 4G; Table S3G – I). The young LINE-2 and SINE-MIR TEs were involved in non-coding and coding RNA processing, metabolism, and basic cellular processes (Fig. 4G; Table S3G – H). Young SINE-Alu TEs overlapping a gene with H3K4me3 loss in sperm were enriched for genes involved in craniofacial and skeletal development (Fig. 4G; Table S3I). We next intersected peaks with H3K4me3 gain or loss to putative sperm and fetal enhancers (Fig. 4H). Peaks with gains in H3K4me3 were solely overrepresented at putative sperm enhancers whereas peaks with H3K4me3 loss were enriched across a broad number of putative developmental enhancers (Fig. 4H). In line with our studies in mice showing that altered sperm H3K4me3 is associated to birth defects in the offspring (Lismer et al., 2021), this analysis suggests that DDT exposures associated with alterations in H3K4me3 may be linked to the increased rates of disease and birth defects in DDT-exposed regions (Bornman et al., 2016; Boucher et al., 2012; Després et al., 2005; Eskenazi et al., 2018).

### *p,p’-*DDE exposure is associated with altered H3K4me3 in sperm at regions that overlap pre-implantation embryo H3K4me3

To assess the possible involvement of sperm H3K4me3 in epigenetic transmission of paternal environmental exposure to DDT we set out to identify whether sperm H3K4me3 peaks were predicted to persist in the early embryo. Regions with H3K4me3 enrichment in the pre-implantation embryo were identified using an existing dataset (Xia *et al*., 2019) and intersected sperm H3K4me3 peaks to pre-implantation embryo H3K4me3 peaks (Fig. S5A – F). A high degree of overlap was identified with 23,582 sperm-to-4-cell persistent H3K4me3 peaks and 7,004 sperm-to-ICM persistent H3K4me3 peaks (Fig. S5G – I; Table S4G – I). We next determined whether peaks with *p,p’*-DDE associated deH3K4me3 retained H3K4me3 in embryos at the 4-cell, 8-cell, and blastocyst stage (Fig. 4I and J, Fig. S5J – M). Interestingly, peaks gaining H3K4me3 in the sperm of DDT exposed men showed a lack of overlap with embryonic H3K4me3 (Fig. 4I, Fig. S5J – K). In contrast, peaks losing H3K4me3 in sperm were highly enriched in H3K4me3 across the studied stages of pre-implantation embryo development (Fisher test p < 0.0001; Fig. 4J, Fig. S5L – M). Revealing a potential cooperativity between DNAme and H3K4me3 in epigenetic transmission of environmental exposures to DDT, peaks with H3K4me3 loss were also highly enriched for dynamic and high DNAme sperm-to-zygote persistent regions (z-score > 0, p < 0.0001, n = 10,000 permutations; Fig. S5N).

Finally, we established whether sperm deH3K4me3 peaks that retained H3K4me3 in the embryo, could potentially impact embryonic gene expression. Promoters bearing H3K4me3 in both sperm and pre-implantation embryos were associated with gene expression at the respective stage of embryogenesis (Fisher test p < 0.0001; see Methods; Fig. S6A – J). Interestingly, most promoters with H3K4me3 loss in sperm have been shown to retain H3K4me3 enrichment in the pre-implantation embryo at genes that are expressed in 4-cell, 8-cell, and blastocyst embryos (Fisher test p < 0.0001; Fig. 4K – M). To assess the functionality of the associated genes, we performed a gene ontology analysis on promoters that retained H3K4me3 in sperm and embryos and overlapped them to genes that were expressed at the corresponding embryonic developmental stage (Fig. S6K – M; Table S3J – L). Significant gene ontology terms included pathways that were critical for development such as placenta vascularization, mRNA processing, chromatin modification, and signalling (Fig. S6K – M; Table S3J – L). Genes of interest included the *BRD1* (chromatin interacting protein), *MAP2K1* (linked to placenta vascularisation), and the paternally expressed imprinted gene *PEG3*. Taken together, our data shows that like DNAme, sperm H3K4me3 may be cumulatively eroded by DDT exposure, and that peaks with H3K4me3 loss at young transposable elements, may escape epigenetic reprogramming after fertilization and lead to deregulated gene expression in the pre-implantation embryo.

### Regions with *p,p’-*DDE associated differential DNAme coincide with deH3K4me3 peaks in sperm

We recently identified regions in sperm bearing both DNAme and H3K4me3, that are involved in important developmental processes (Lambrot *et al*., 2021). We consequently wanted to determine if DMRs and deH3K4me3 co-occurred at the same genomic loci. From the 830,188 MCC-seq target regions, 157,753 intersected H3K4me3 peaks (Fig. 5A). Conversely, 34,741 H3K4me3 peaks overlapped MCC-seq target regions (Fig. 5A), which can be explained by the preferential distribution of H3K4me3 peaks in CpG enriched regions which are well covered in the capture design (Chan et al., 2019). Interestingly, 1,744 DMRs overlapped a H3K4me3 peak and 106 DMRs overlapped a deH3K4me3 peak in South African sperm (Fig. 5A). DMRs that gained DNAme but lost deH3K4me3 (= 88 DMRs) showed the highest overlap (Fig. 5B). DNAme gain DMRs and H3K4me3 loss regions may be the most biologically relevant in the embryo because based on our findings, they 1) may conserve DNAme and / or H3K4me3 in the embryo and 2) were more functionally relevant for development (Fig. 2 and 4). Overlapping DNAme gain DMRs and H3K4me3 loss regions were more likely found at < 5 kb to the TSS at CpG shores compared to the other differential regions without both H3K4me3 loss and DNAme gain (Fig. 5C – D; z-score > 0; p < 0.0001; n = 10,000 permutations). These overlapping regions were also more likely enriched at H3K4me3 sperm-to-4-cell and sperm-to-ICM persistent regions, SINE-MIRc, hESC enhancers, as well as low and dynamic DNAme persistent regions than other differential regions with only one change in either DNAme or H3K4me3 (Fig. 5E; z-score > 0; p < 0.0001; n = 10,000 permutations). Gene ontology analysis of peaks with a loss of H3K4me3 and a gain in DNAme revealed enrichment for genes involved in reproduction, spermatogenesis, embryo development and neurodevelopment (Fig. 5F and Table S3M). Representative visualizations of dose response differences in enrichment for DNAme and H3K4me3 in overlapping regions are shown for HERV/LTR43 and the gene TRAPPC9 (Fig. 5G), which has been implicated in cognitive disability and microcephaly (Mochida et al., 2009). Interestingly, a search for regulatory elements in HERV/LTR43 (Ito et al., 2017), revealed that it is bound by numerous transcription factors implicated in development and disease. These included PAX5 (implicated in the developing CNS and testis), FOS (regulator of cell proliferation and differentiation), MYC (proto-onogene), NFKB1(inflammatory regulator) and NFE2 (blood vessel and bone development).

**Fig. 5:**
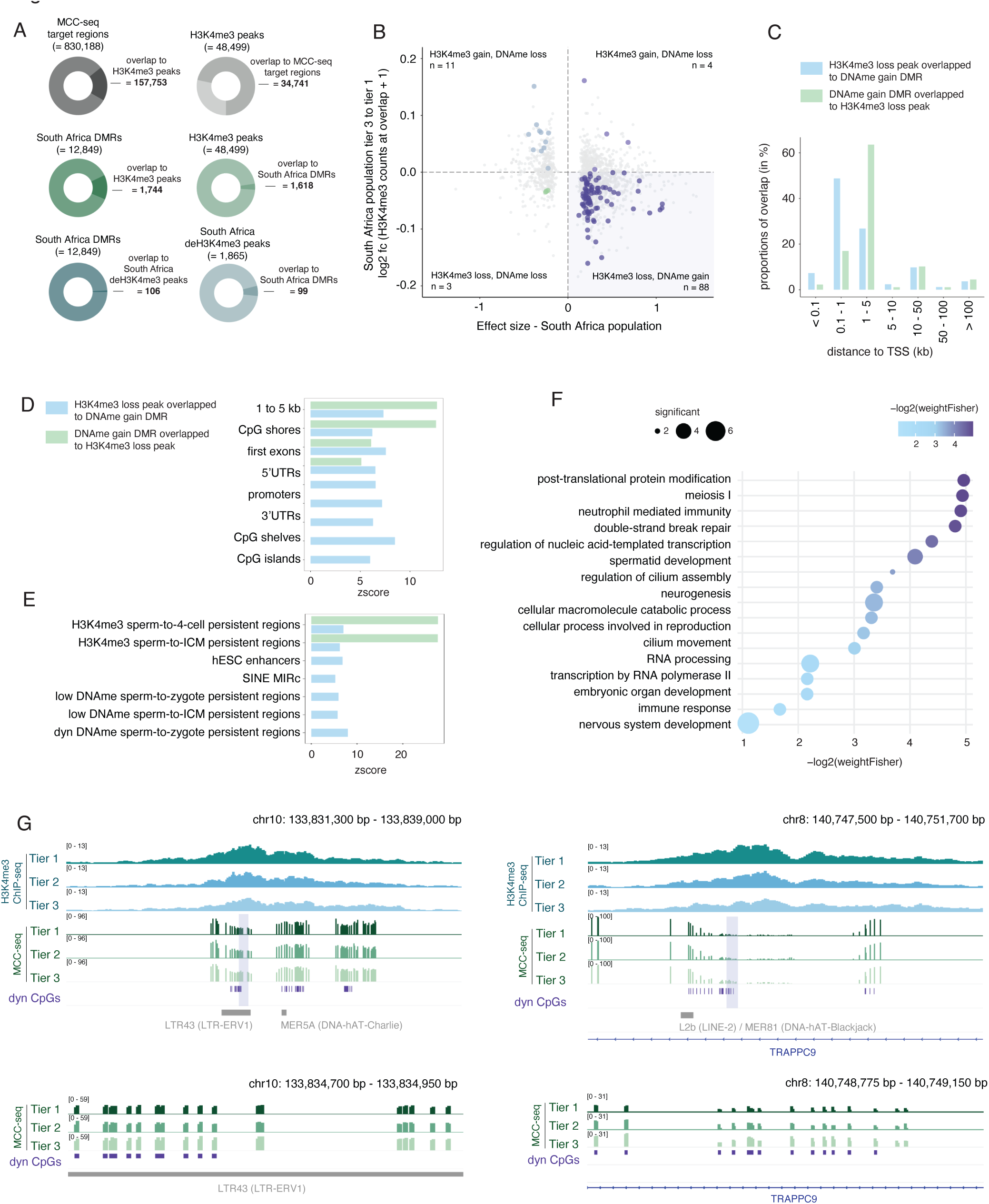
deH3K4me3 peaks and DMRs intersect in sperm of *p,p’-*DDE -exposed South African men. (A) Proportion of MCC-seq target regions that overlap a H3K4me3 peak in sperm (overlap = 157,753; dark grey donut plot), proportion of H3K4me3 peaks in sperm that overlap a MCC-seq target region (overlap = 34,741; light grey donut plot), proportion of South Africa DMRs that overlap a H3K4me3 peak in sperm (overlap = 1,744; dark green donut plot), proportion of H3K4me3 peaks in sperm that intersect a South Africa DMR (overlap = 1,618; light green), proportion of South Africa DMRs that overlap a deH3K4me3 peak in sperm (overlap = 106; dark blue), proportion of South Africa deH3K4me3 peaks in sperm that intersect a DMR in sperm (overlap = 99; light blue). (B) Scatter plot corresponding to South African population tier 3 to tier 1 log2 fold change (H3K4me3 counts at overlapping DMRs + 1) relative to South Africa beta value at the DMRs. Grey dots correspond to DMRs that do not overlap with a deH3K4me3 peak. DMRs intersecting a deH3K4me3 peak are denoted by colour. H3K4me3 loss + DNAme gain overlap was the predominant overlap (n = 88 out of 106). (C) Distribution of peaks with H3K4me3 loss overlapped to DNAme gain DMR (blue) and DNAme gain DMR overlapped to H3K4me3 loss region (green) relative to the TSS. (D) Enrichment for peaks with H3K4me3 loss overlapped to DNAme gain DMR (blue) and DNAme gain DMR overlapped to H3K4me3 loss region (green) at genic and CpG annotations. Positive enrichments are determined by Z scores using the Bioconductor package regioneR. For all annotations displayed, p < 0.0001 and n = 10,000 permutations of random regions resampled from deH3K4me3 peaks (blue) and DMRs (green), respectively. (E) Enrichment for peaks with H3K4me3 loss overlapped to DNAme gain DMR (blue) and DNAme gain DMR overlapped to H3K4me3 loss region (green) at transposable element annotations and characterized DNAme / H3K4me3 persistent regions (see Fig. S2 and S5). Positive enrichments are determined by Z scores using the Bioconductor package regioneR. For all annotations displayed, p < 0.0001 and n = 10,000 permutations of random regions resampled from deH3K4me3 peaks (blue) and DMRs (green), respectively. (F) Selected significant pathways from gene ontology analysis on peak with H3K4me3 loss that overlaps a DNAme gain DMR at promoters and 1 - 5 kb genic space. Size of dots corresponds to the number of significant genes in the pathway. Colour and position of dots correspond to -log2(weightFisher). (G) Representative IGV tracks of peak with H3K4me3 loss and DNAme gain in South African sperm at an LTR-ERV1 and in the TRAPPC9 genic space. Purple shaded box corresponds to the DMR. Tracks below are a zoom at the DMR. Dynamic CpGs are indicated in purple.

## Discussion

There is now an abundance of alarming evidence from experimental models that paternal exposures can negatively impact offspring phenotypes (reviewed in Skvortsova et al., 2018). In rodent models of paternal epigenetic inheritance DNAme, chromatin and non-coding RNA have been implicated in response to environmental challenges including diet (Chen et al., 2016; Lismer *et al*., 2021; Ly et al., 2017; Pepin *et al*., 2022; Sharma et al., 2016), trauma (Gapp et al., 2020) and toxicants (Anway and Skinner, 2006; Maurice *et al*., 2021; Skinner *et al*., 2018). In humans, the intergenerational effects on the health of children as a consequence of maternal exposures to environmental factors including toxicants are well studied. In contrast, few studies have examined paternal modes of transmission that impact child health, and these have been predominantly epidemiology based (reviewed in Soubry et al., 2014). A limitation of epigenetic inheritance studies is the difficulty to go beyond association of an epigenetic alteration from an exposure, and a change of function in the offspring. This has been particularly challenging in human studies where it is difficult to access tissues in the next generation that are matched to a father, as was the case in this study. Here, we addressed this limitation by combining the findings from the sequencing data generated in this study, which identified environmentally responsive sites of sperm DNA methylation and H3K4me3, to mined epigenomic and gene expression data from prior studies that used human embryos (Guo *et al*., 2014; Xia *et al*., 2019). This approach allowed us to advance our understanding of paternal epigenetic inheritance by identifying potential regions of environmental sensitivity in sperm that could be further studied to confirm embryonic transmission and functions in the next generation.

Other important limitations in the presented study included the relatively small sample size. Unlike samples acquired through most clinical studies, these samples were highly precious and unique coming from remote areas of the world from small indigenous populations. The size and regions of the populations limit the metadata that can be publicly released due to risk of participant identification (General Data Protection Regulations). Because the Greenlandic cohort samples were from the INUENDO biobank, it was not possible to coordinate factors with the prospective South African study. As such, the populations differed in the timing of collection, in mean age (25 versus 31 years of age in South African and Greenlandic men respectively) and the site of *p,p’-*DDE measurement. Moreover, the Greenlandic Inuit men were proven fertile whereas the fertility status of the South African men was unknown. Semen analysis has been performed on both populations in prior studies. There was no association with *p,p’-*DDE and sperm concentration or morphology, while higher doses were associated with reduced motility in samples from Greenlandic Inuit (Toft et al., 2006). Likely reflecting higher exposures in the South African population in a previous study we identified increased incidences of teratozoospermia, asthenozoospermia and lower motility (Aneck-Hahn et al., 2007). In this study, to best match the fertility status of the Greenlandic Inuit men, we selected South African samples with normal sperm counts and all but one sample had normal motility. It was not deemed possible for socio-cultural reasons to request participants to provide information on fertility status. Altered fertility status has been associated with changes in DNAme, yet the identification of CpGs that are consistently changed in men with infertility linked to poor embryo quality is inconclusive (Aston et al., 2015). Conceivably some of the DNAme differences detected between the two populations may be linked to fertility status. However, given that there is no clear infertile DNAme signature we cannot for example remove a designated set of CpGs from our analysis. Consideration must also be given to the impact of unidentified environmental confounders such as exposures to other EDC compounds and an inability to discriminate between acute versus long-term exposures. Additionally, we do not know how stable the sperm epigenome is and whether *p,p’-*DDE associated alterations are temporary or permanent. Large-scale intergenerational studies that include sperm epigenome analyses are needed to better understand the connections between paternal exposures, the sperm epigenome, and child development. However due to cultural and societal differences such studies are inherently challenging in the context of these population and more suited to a clinical setting.

Notwithstanding the population differences mentioned above and the range in *p,p’-*DDE body burdens, there was a high degree of consistency of *p,p’-*DDE associated DNAme alterations. While we cannot rule out selection bias or residual confounding factors, the overlapping DNAme changes confirmed in two independent populations of different selection / confounder structure, clearly indicates that this overlap is unlikely to be driven by bias. In both populations, DMRs localized to genes and regulatory elements that reflect the increased susceptibility to disease and developmental disorders associated with EDC and DDT exposures, including increased birth defects and neurodevelopmental impairments (Alan S. Brown et al., 2018; Caporale et al., 2022; de Jager et al., 2009; Van Oostdam *et al*., 2005). Strikingly, 17.5% of regions with altered DNAme in Greenlandic sperm were also altered in South African sperm. DNAme was preferentially altered in the intergenic space, at dynamic DNAme CpGs where there were more gains than losses of DNAme. The intergenic space is enriched for TEs including retroviruses which are controlled by DNA and histone methylation (Goodier, 2016; Liu et al., 2014). TEs function as promoters, transcription factor binding sites and enhancers implicated in pluripotency (Senft and Macfarlan, 2021). Enhancer-promoter interactions are facilitated by the CTCF binding sites in TEs which in turn influence chromatin looping (Diehl et al., 2020). Factors such as aging, infection, or hormones have been postulated to reactivate human endogenous retroviruses (ERVs) (Küry et al., 2018). Perhaps reflecting the endocrine disrupting activity of DDT metabolites, was the specific sensitivity associated with DDT exposure to the DNAme changes observed in the long terminal repeat ERVs (LTR-ERV1, LTR-ERVK, LTR-ERVL). Expression of ERV families is temporally and spatially regulated during human embryogenesis, and function in gene regulation as alternate promoters and enhancers (Göke et al., 2015; Rodriguez-Terrones and Torres-Padilla, 2018). Likewise, deH3K4me3 was over-represented at LINE-2, SINE-MIR, and SINE-Alu, while LTR-ERV1, overlapped both deH3K4me3 and DMRs. These epigenetic changes at TEs may in turn serve in the epigenetic inheritance of DDT-associated phenotypes. TEs known to be resistant to epigenetic reprogramming in human pre-implantation embryos include ERVs, SINEs and LINEs (Guo *et al*., 2014; Zeng et al., 2022). All factors taken together, this study bolsters the evidence that in humans like plants and animals (Heard and Martienssen, 2014), TEs are attractive genome mediators of environmentally influenced non-genetic inheritance. What remains unknown for all species is how epigenetic marks at TEs escape embryonic reprogramming.

*p,p’-*DDE may influence epigenetic programming linked to TEs, and epigenetic inheritance in sperm via its endocrine disruptor effects on androgenic and estrogenic signaling. Androgen regulates genes in one carbon metabolism including glycine *N*-methyltransferase, cystathionine β-synthase and ornithine decarboxylase and in this way can impact methylation of DNA and histones (Corbin and Ruiz-Echevarría, 2016). While direct evidence is lacking, one study connected the EDC bisphenol A to epigenetic changes that were mitigated by folate supplementation. Agouti (A*^vy^*) mice that have metastable epiallele determining coat color were exposed to BPA and consequently had offspring with a coat color change. This change corresponded to methylation changes at the TE upstream of the A*^vy^*. This BPA induced phenotype was prevented in the presence of folic acid (Dolinoy et al., 2007). Therefore, it is plausible that exposure of men *p,p’-*DDE leads to the altered sperm epigenome through similar disruption of one carbon metabolism, to negatively alter the sperm epigenome.

We identified regions in sperm bearing altered DNAme and H3K4me3 in association with DDT and *p,p’-*DDE exposure that were predicted to persist in the embryo. This is a plausible route for paternally mediated epigenetic inheritance that leads to effects in offspring and is supported by exposure studies in animal models (Anway and Skinner, 2006; Lismer *et al*., 2021; Ly *et al*., 2017; Maurice *et al*., 2021; Oluwayiose et al., 2021; Skinner et al., 2013). Of relevance are two studies of paternal DDT exposures that were associated with transgenerational alterations in sperm DNAme profiles, obesity and developmental abnormalities (Maurice *et al*., 2021; Skinner *et al*., 2013). However, in humans the connection between paternal exposures, the sperm epigenome and health in the next generation is underexplored. In fact, there are few large-scale epidemiological studies that suggest paternal exposures effect health in children except for several exploring links to childhood cancers. For example, paternal exposure to herbicides was associated with a significant risk for astrocytoma, a childhood brain cancer (Shim et al., 2009). Similarly, paternal exposure to pesticides was associated with higher risk for childhood acute myeloid leukemia (Patel et al., 2020). We did not find clear epigenomic connections suggesting a link between DDT and *p,p’-*DDE exposure to childhood cancer. However, a consistent finding for *p,p’-*DDE -associated DMRs and deH3K4me3 peaks in sperm was their occurrence at genes or regulatory elements related to neurodevelopment and neurofunction. The health connections between paternal DDT and *p,p’-*DDE exposures and neurodevelopment are unknown. The incidence of neurodevelopmental impacts associated with maternal serum *p,p’-*DDE in Greenlandic children of the INUENDO cohort have been reported. Pre- natal and post-natal exposures via the mother to *p,p’-*DDE and PCB-153 was associated with an increased incidence of hyperactivity in children (Rosenquist Aske et al., 2017). Interestingly, maternal exposures to EDCs have been associated with reductions in cognitive function such as delayed language acquisition. For example, a Finnish birth cohort study of > 1 million mother-child pairs found that the risk of childhood autism was significantly increased with serum *p,p’-*DDE levels that were in the highest 75^th^ percentile (Alan S. Brown *et al*., 2018). As in most birth cohorts the paternal contribution was not studied, yet conceivably paternal *p,p’-*DDE exposure may also have contributed to the findings of increased autism risk. Clearly there is a gap in epidemiological evidence relating paternal toxicant exposures to adverse effects in children. Our inability to track childhood health in relation to paternal exposures in the studied populations prevented us from making these essential connections.

Despite a multitude of documented adverse health effects attributed to DDT exposure, its effectiveness as an anti- malarial control method has led to its continued use in 14 countries (van den Berg, 2010). The impacts of DDT on human health are not only restricted to regions of use since DDT is transported over long distances in the environment by weather patterns and ocean currents (Simonich Staci and Hites Ronald, 1995). Due to climate change, the long-range contamination, environmental concentration, persistence, and bioaccumulation of DDT and its persistent transformation product, *p,p’-*DDE, are predicted to worsen (Teran et al., 2012). Human exposures to DDT are consequently expected to increase in regions distant from its use (Teran et al., 2012). Taking this into consideration with the implication that DDT exposure alters the sperm epigenome and may be implicated in epigenetic inheritance of disease, it is essential to consider alternative methods for malarial control. Additionally, our findings highlight the need for future regulatory decision-making to be based on chemical risk assessments that incorporate experimental toxicology and epigenetic profiling of sperm.

## Acknowledgements

We thank Drs. Olusola Sotunde and Hope Weiler for their assistance with the study, as well as the South African Medical Research Council for funding (SIR grant), the fieldworkers, and participants from the different villages in Vhembe district, Limpopo province, South Africa. We would like thank Genome Quebec for the construction of the WGBS libraries. We thank Dr. Guillaume Bourque from the McGill Genome Centre for his support for the MCC-Seq project and Marie-Michelle Simon for preparing the MCC-Seq libraries. Epigenomic sequencing was performed at the McGill Genome Center under the guidance of Dr. Tony Kwan. We appreciate advice from Dr. Yann Joly (McGill, Research Director of the Centre of Genomics and Policy) regarding regulatory ethics related to genomic data access. This research was enabled in part by support provided by Calcul Quebec and the Digital Research Alliance of Canada.

## Author contributions

**A. Lismer:** methodology, data curation, investigation, formal analysis, visualization, writing – original draft, editing / reviewing

**X. Shao:** software, methodology, formal analysis, writing – editing / reviewing

**M.C. Dumargne:** methodology, investigation, formal analysis

**C. Lafleur**: methodology, investigation

**R. Lambrot**: methodology, investigation

**D. Chan**: formal analysis, visualization, writing – editing / reviewing

**G. Toft** data curation, writing – editing / reviewing

**J.P. Bonde:** data curation, writing – editing / reviewing

**A.J. MacFarlane:** methodology, investigation, funding acquisition, writing – editing / reviewing

**M. Bornman:** methodology, investigation, data curation, writing – editing / reviewing

**N. Aneck-Hahn:** methodology, investigation, data curation, writing – editing / reviewing, project administration

**S. Patrick:** methodology, investigation, data curation, writing – editing / reviewing, project administration

**J.M. Bailey:** conceptualization, funding acquisition

**C. de Jager:** conceptualization, methodology, funding acquisition, writing – editing / reviewing

**J.M. Trasler:** supervision, conceptualization, methodology, funding acquisition, writing – editing / reviewing

**V. Dumeaux**: software, formal analysis, visualization, writing – editing / reviewing

**S. Kimmins**: supervision, conceptualization, methodology, data curation, funding acquisition, writing – original draft, editing / reviewing, project administration

## Data and materials availability

Compliance with General Data Protection Regulations in Denmark permits access to the Greenlandic Inuit MCC-seq data upon request via material transfer agreement only. South African phenotypic and epigenomic data will be made available by request through the European Genome-Phenome Archive (EGA) upon publication. Further information about EGA can be found on https://ega-archive.org (Freeberg et al., 2022).

## Supplemental Tables

**Table S1:** Aggregated participant information from Greenlandic Inuit and South African VhaVenda cohorts.

**Table S2:** MCC-seq read statistics, differentially methylated CpGs, hotspots, and differentially methylated regions, in Greenlandic Inuit and South African VhaVenda sperm.

**Table S3:** Gene ontology analyses on differentially methylated regions and differentially enriched H3K4me3 peaks in Greenlandic Inuit and South African VhaVenda sperm.

**Table S4:** Sperm-to-embryo predicted persistent DNAme and H3K4me3 regions.

**Table S5:** ChIP-seq read statistics and differentially enriched H3K4me3 peaks in South African VhaVenda sperm.

---

**Fig. S1:**
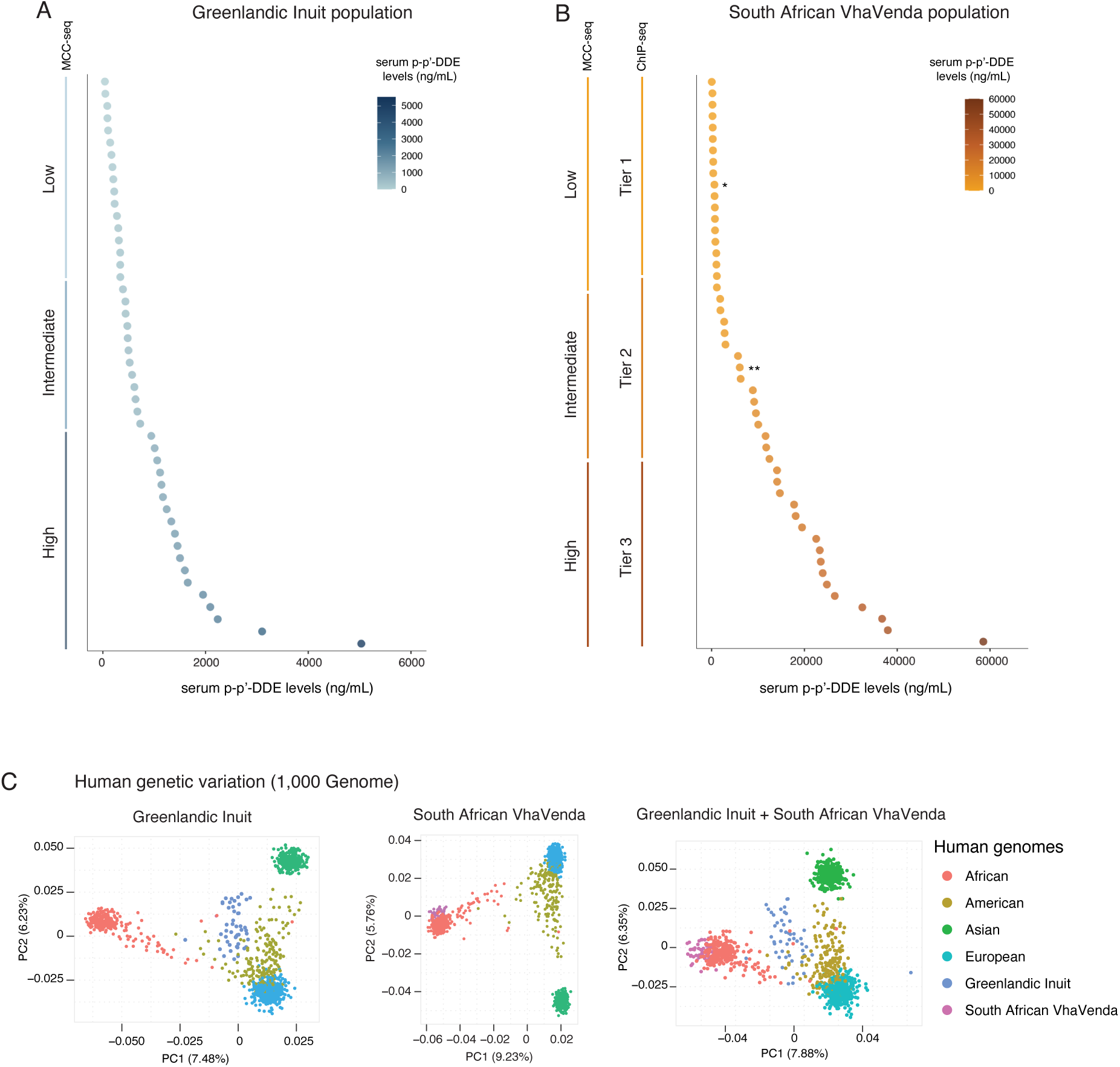
Characterization of Greenlandic and South African sperm relative to serum *p,p*’*-*DDE levels and single- nucleotide polymorphisms. (A) Distribution of *p,p’*-DDE serum levels in Greenland men (n = 48) from this study. (B) Distribution of *p,p’*-DDE serum levels in South African men (n = 51) from this study. Exposure tiers for H3K4me3 analysis are indicated on the y-axis. (*) denotes sample with only an MCC-seq dataset and (**) denotes sample with only a ChIP-seq dataset. (C) Principal component analysis plot on genotype profiles of Greenland and South African populations. Chromosome 1 genotype data from 1,000 genomes were used as the reference genotype profile (the human population genetic background) to compare with Greenland (indigo) and South African cohorts (magenta).

**Fig. S2:**
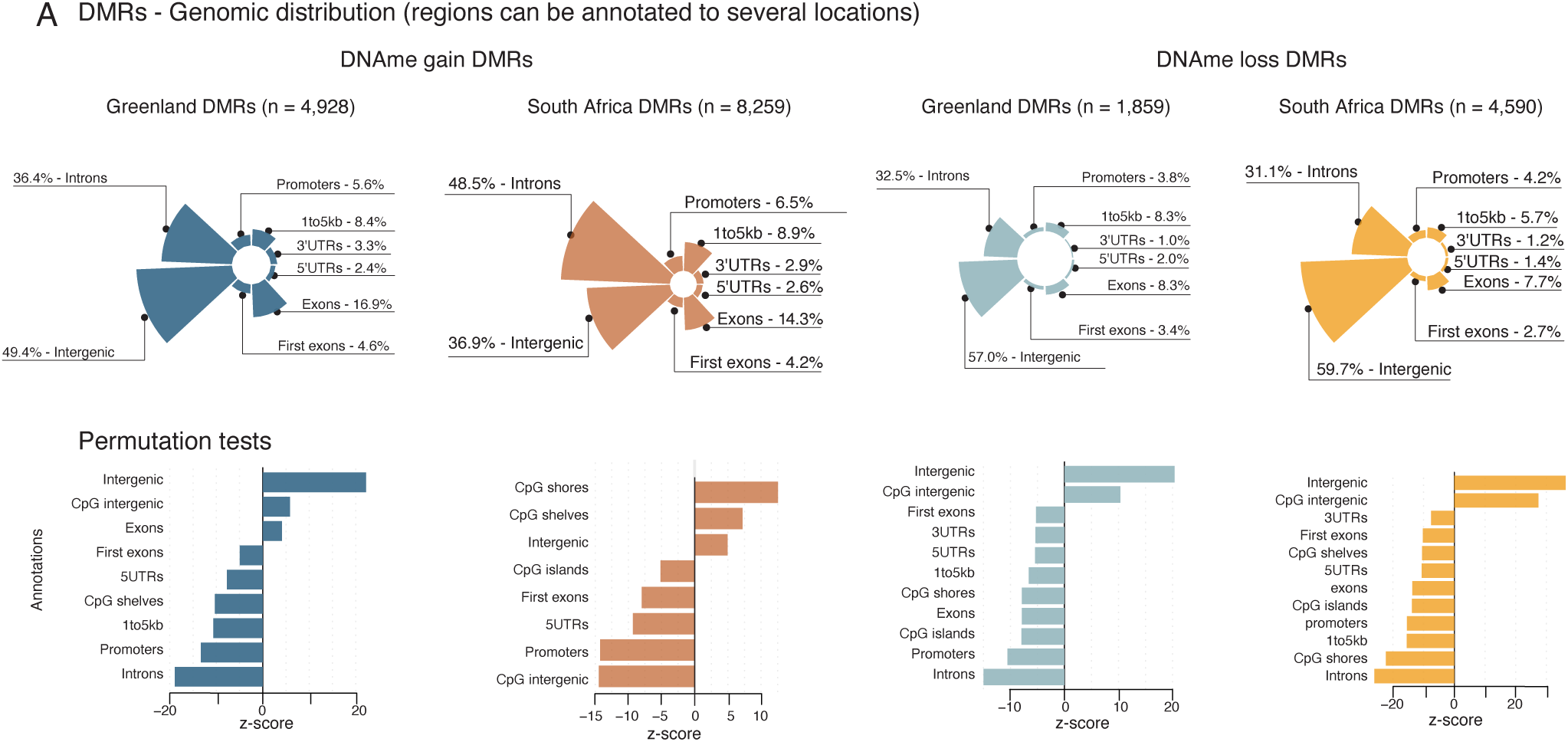
Genic and transposable element characterization of differentially methylated regions. (A) Genic distributions and genic / CpG enrichments at DNAme gain or loss DMRs in Greenland or South Africa sperm. Positive enrichments are determined by Z scores using the Bioconductor package regioneR. For all annotations displayed, p < 0.0001 and n = 10,000 permutations of random regions (of the same size) resampled from the targeted MCC-seq regions.

**Fig. S3:**
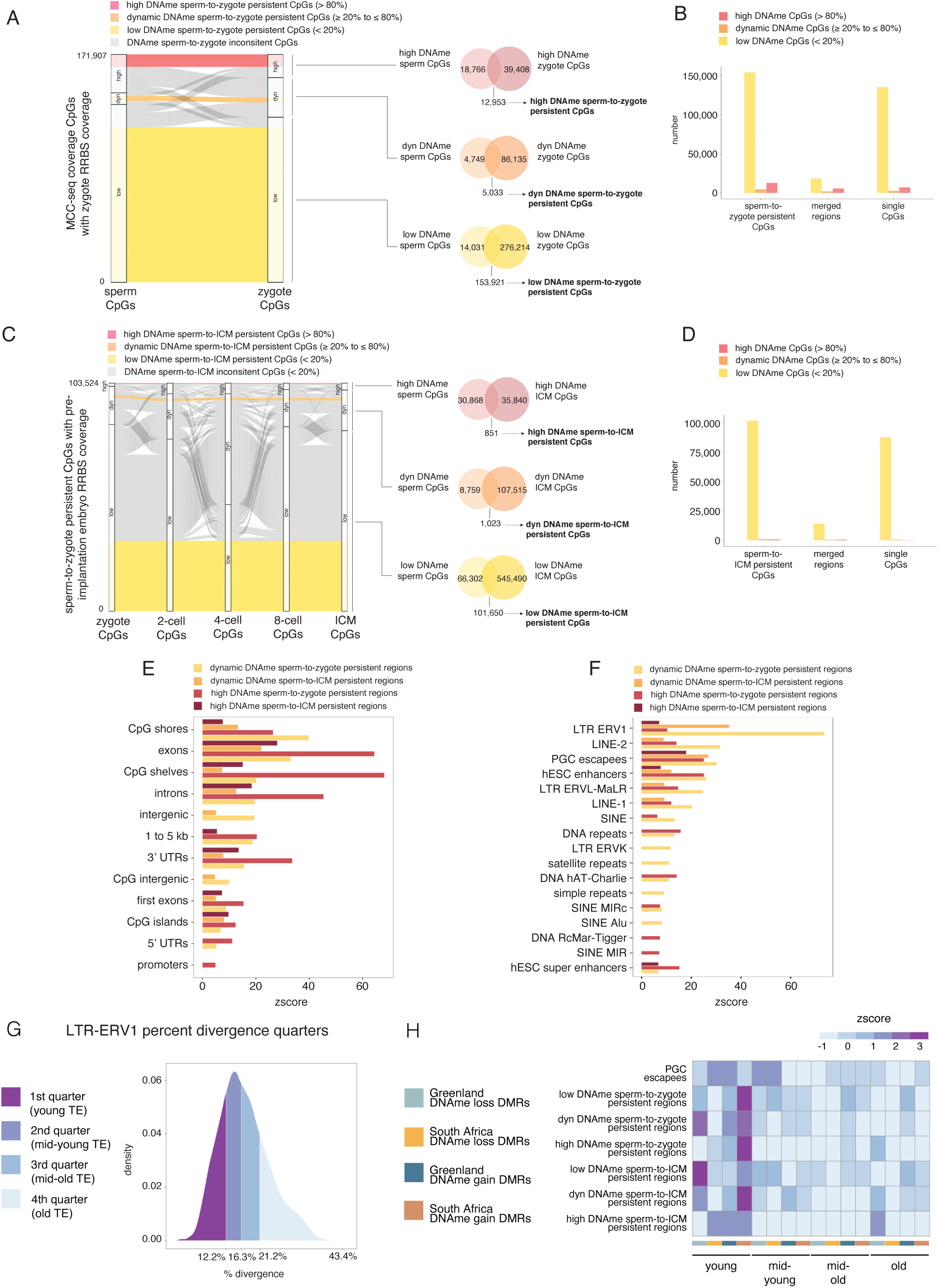
Identification of predicted persistent DNAme regions from sperm to the pre-implantation embryo. (A) Alluvial plot identifying CpGs that retain the same level of DNAme from sperm to the zygote. High DNAme sperm to zygote persistent CpGs are shown by red ribbon, dynamic DNAme sperm to zygote persistent CpGs are denoted by orange ribbon, low DNAme sperm-to-zygote persistent CpGs are characterized by yellow ribbon. Grey ribbon corresponds to non-persistent DNAme CpGs. Each node indicates the DNAme level for the specific cell type. Y-axis corresponds to MCC-seq coverage CpGs that have zygote RRBS coverage. Venn diagrams show sperm-to-ICM persistent CpGs for high, dynamic, or low DNAme levels. (B) Number of total DNAme sperm-to-zygote persistent CpGs, DNAme sperm-to-zygote persistent regions, and single DNAme sperm-to-zygote persistent CpGs, for low DNAme (yellow), dynamic DNAme (orange), and high DNAme (red). DNAme sperm-to-zygote persistent regions were called by merging DNAme sperm-to-zygote persistent CpGs separated by a maximum distance of 500 bp. (C) Alluvial plot identifying CpGs that retain the same level of DNAme across all stages of pre-implantation embryogenesis. High DNAme sperm to zygote persistent CpGs are shown by red ribbon, dynamic DNAme sperm to zygote persistent CpGs are denoted by orange ribbon, low DNAme sperm-to-zygote persistent CpGs are characterized by yellow ribbon. Grey ribbon corresponds to non-persistent DNAme CpGs. Each node indicates the DNAme level for the specific cell type. Y-axis corresponds to sperm-to-zygote persistent CpGs that have RRBS coverage across the studied stages of pre-implantation embryogenesis. Venn diagrams show sperm-to-ICM persistent CpGs for high, dynamic, or low DNAme levels. (D) Number of total DNAme sperm-to-ICM persistent CpGs, DNAme sperm-to-ICM persistent regions, and single DNAme sperm-to-ICM persistent CpGs, for low DNAme (yellow), dynamic DNAme (orange), and high DNAme (red). DNAme sperm-to-ICM persistent regions were called by merging DNAme sperm-to-ICM persistent CpGs separated by a maximum distance of 500 bp. (E) Enrichment for characterized DNAme persistent regions at genic annotations. Positive enrichments are determined by Z scores using the Bioconductor package regioneR. For all annotations displayed, p < 0.0001 and n = 10,000 permutations of random regions (of the same size) resampled from the targeted MCC-seq regions. (F) Enrichment for characterized DNAme persistent regions at transposable elements (RepeatMasker hg19 library 20140131). Positive enrichments are determined by Z scores using the Bioconductor package regioneR. For all annotations displayed, p < 0.0001 and n = 10,000 permutations of random regions (of the same size) resampled from the targeted MCC-seq regions. (G) Density plot of percent divergences for LTR-ERV1 transposable elements in the hg19 genome. Color shading corresponds to quantile cutoffs and associated age categories. (H) Enrichment of young, mid-young, mid-old, and old LTR-ERV1 transposable elements that overlap a PGC escapee and/or characterized sperm-to-pre-implantation-embryo persistent region (see Fig. S2) at given DMR.

**Fig. S4:**
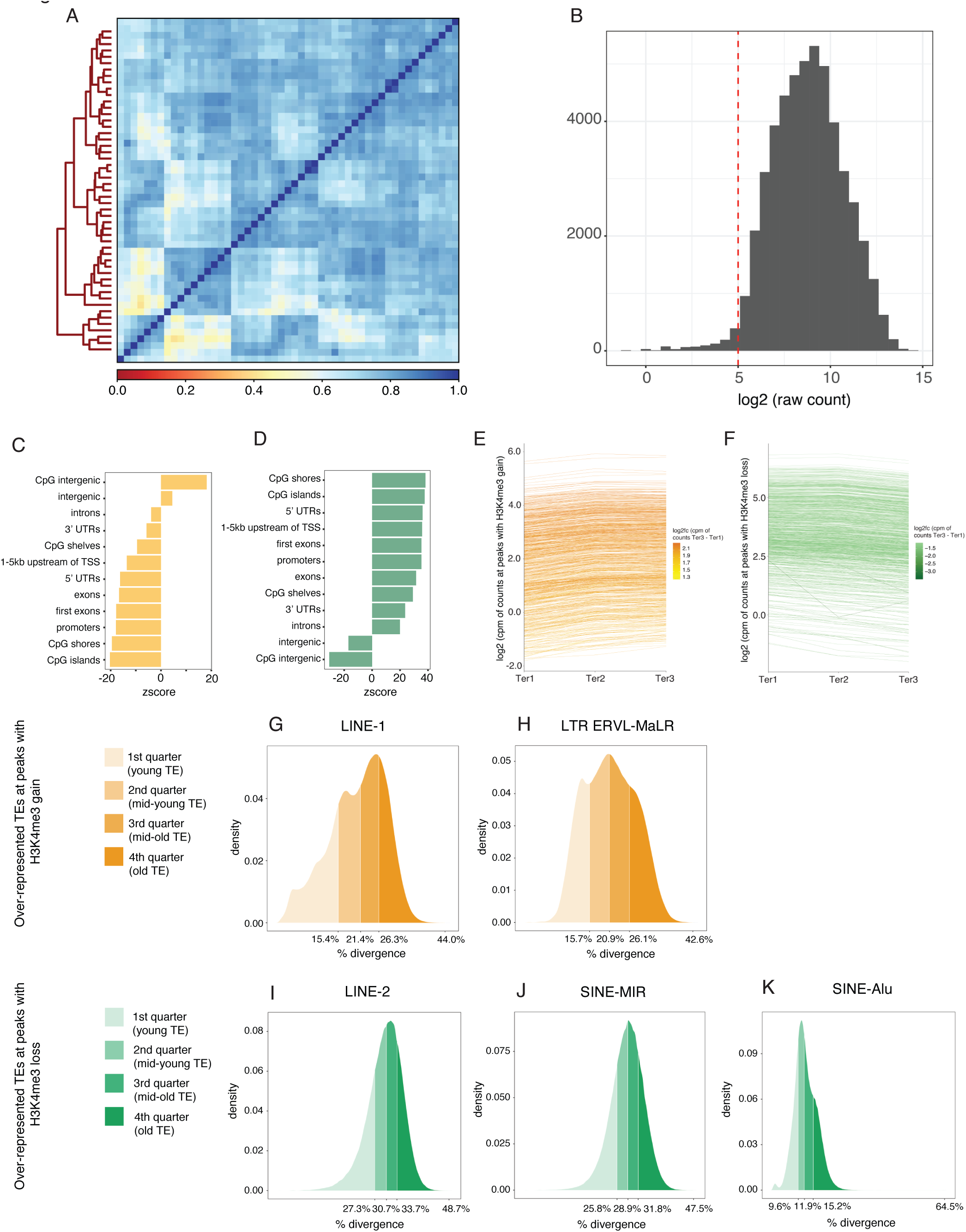
Genic and transposable element characterization of differentially enriched H3K4me3 peaks. (A) Heatmap depicting spearman correlations between samples based on normalized counts of H3K4me3 peaks identified in our reference human dataset (50,117 peaks; REF). Normalized counts are reads per kilobase per million mapped reads. (B) Distribution of the log-transformed median raw counts of H3K4me3 reference peaks across samples in this dataset. We selected peaks with log2 median counts above 5 for downstream analyses (n = 48,499 peaks). (C) Enrichment of peaks with H3K4me3 gain at genic annotations. Positive and negative enrichments are determined by Z scores. For all annotations displayed, p < 0.0001 and n = 10,000 permutations of random regions (of the same size) resampled from sperm H3K4me3 peaks. (D) Enrichment of peaks with H3K4me3 loss at genic annotations. Positive and negative enrichments are determined by Z scores. For all annotations displayed, p < 0.0001 and n = 10,000 permutations of random regions (of the same size) resampled from sperm H3K4me3 peaks. (E) Line diagram for regions with H3K4me3 gain in VhaVenda sperm where each line corresponds to the log2 cpm of H3K4me3 counts at individual regions with H3K4me3 gain in sperm across men from ter1, ter2, and ter3 *p,p’*-DDE exposure levels. (F) Line diagram for regions with H3K4me3 loss in VhaVenda sperm where each line corresponds to the log2 cpm of H3K4me3 counts at individual regions with H3K4me3 loss in sperm across men from ter1, ter2, and ter3 *p,p’*-DDE exposure levels. (G - H) Density plots of percent divergences for LINE-1 (E) and LTR ERVL-MaLR (F) transposable elements in the hg19 genome. Color shading corresponds to quantile cutoffs and associated age categories. (I - K) Density plots of percent divergences for LINE-2 (G) and SINE-MIR (H), and SINE-Alu (I) transposable elements in the hg19 genome. Color shading corresponds to quantile cutoffs and associated age categories.

**Fig. S5:**
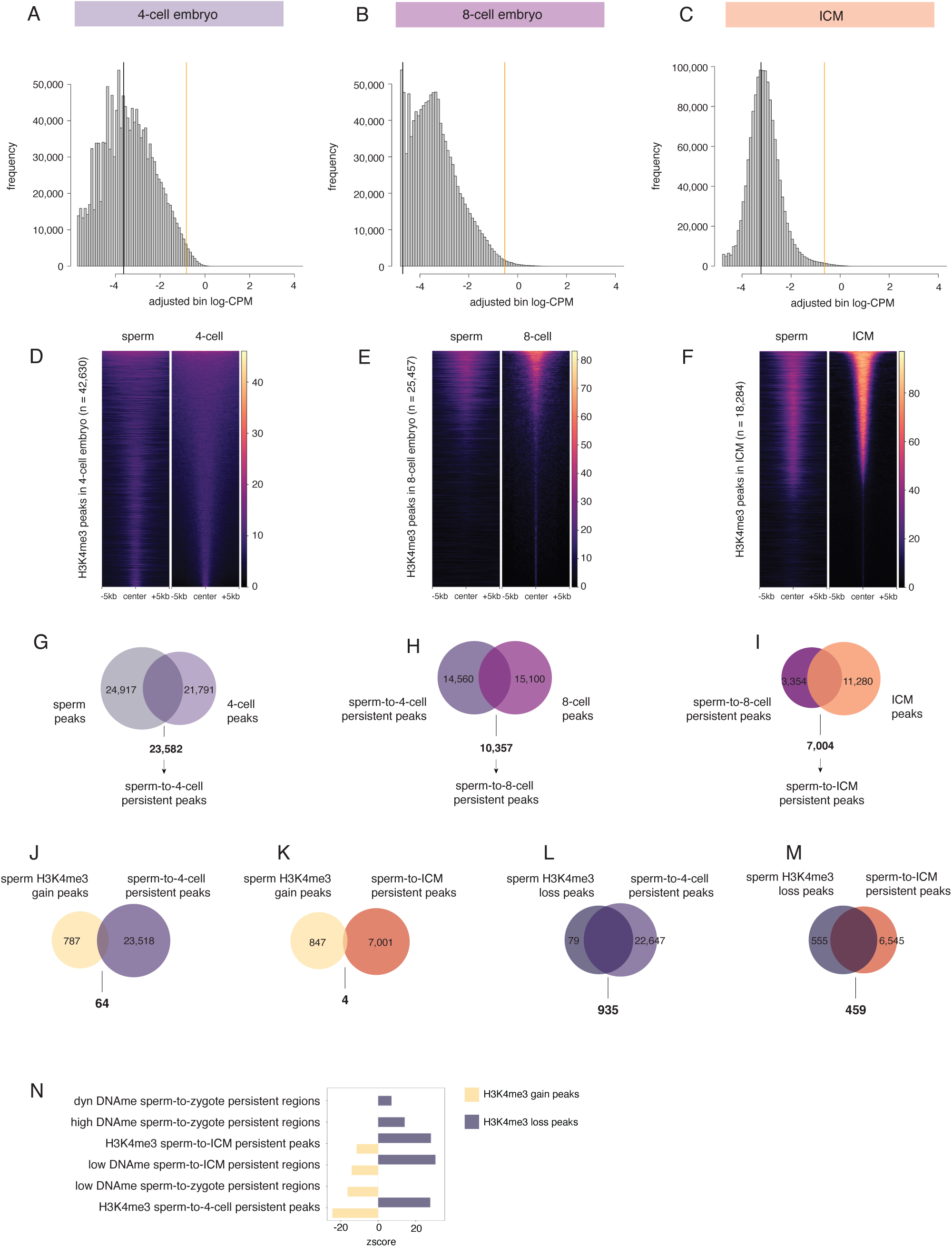
Identification of predicted persistent H3K4me3 peaks from sperm to the pre-implantation embryo. (A - C) Histograms of background read abundance as determined by the number of H3K4me3 ChIP-seq reads in 2000 bp windows tiled across the hg19 genome for 4-cell embryos (A), 8-cell embryos (B), and ICM (C). An abundance threshold was set at ≥ log2(7) fold over background for 4-cell embryos (A), ≥ log2(18) fold over background for 8-cell embryos (B), and ≥ log2(6) fold over background for ICM (C). Windows below this threshold were filtered out for downstream analysis. Remaining windows less than 5,000 bp (A), 6,000 bp (B), or 2,000 bp (C) apart were merged to generate H3K4me3 peaks with a maximum width of 20,000 bp. Peaks identified as enriched for H3K4me3 in the pre-implantation embryo were then inspected and confirmed in IGV. (D - F) H3K4me3 signal intensity enrichment heatmaps at +/- 5kb center of 4-cell (D), 8-cell (E), or ICM (F) peaks relative to sperm or pre-implantation embryo signal, and sorted by pre-implantation embryo H3K4me3 signal intensity. (G - I) Overlap between sperm H3K4me3 peaks and 4-cell H3K4me3 peaks (= 23,582; G), sperm-to-4-cell persistent H3K4me3 peaks and 8-cell H3K4me3 peaks (= 10,357; H), sperm-to-8-cell persistent H3K4me3 peaks and ICM H3K4me3 peaks (= 7,004; I). (J - K) Overlap between sperm H3K4me3 gain peaks and 4-cell H3K4me3 peaks (= 64; J), sperm H3K4me3 gain peaks and sperm-to-ICM persistent H3K4me3 peaks (= 4; K). (L - M) Overlap between sperm H3K4me3 loss peaks and 4-cell H3K4me3 peaks (= 935; L), sperm H3K4me3 loss peaks and sperm-to-ICM persistent H3K4me3 peaks (= 459; M). (N) Enrichment for peaks with H3K4me3 gain or loss at characterized DNAme / H3K4me3 persistent regions (see Fig. S2 and S5). Positive enrichments are determined by Z scores using the Bioconductor package regioneR. For all annotations displayed, p < 0.0001 and n = 10,000 permutations of random regions (of the same size) resampled from sperm H3K4me3 peaks.

**Fig. S6:**
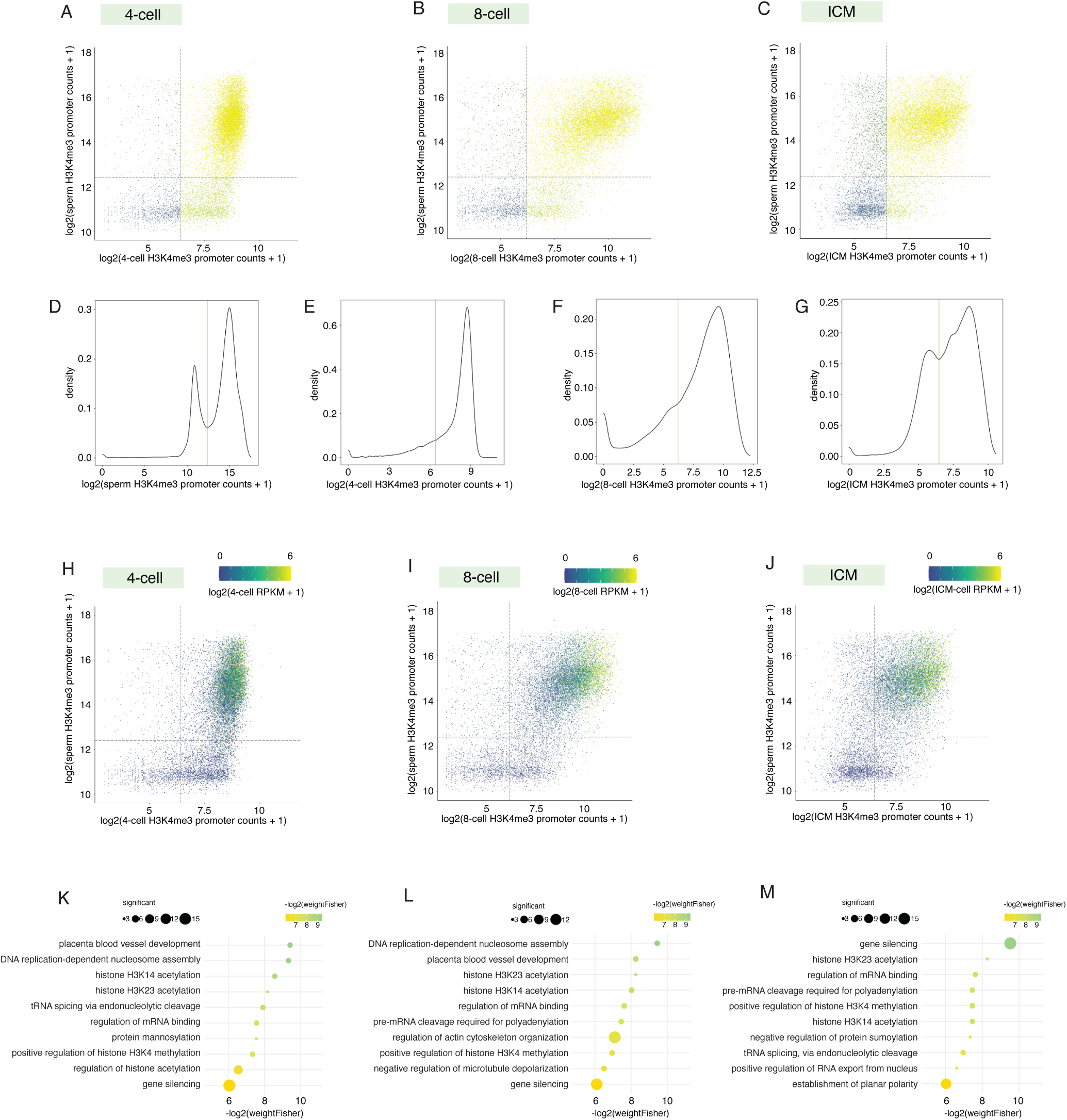
Promoters marked by H3K4me3 in sperm and pre-implantation embryos correspond to expressed genes in the embryo. (A - C) Scatterplots where the x axis corresponds to the log2 (pre-implantation embryo H3K4me3 promoter counts + 1) and the y axis corresponds to the log2 (sperm H3K4me3 promoter counts + 1) at +/- 1 kb TSS of the hg19 genome. Colour of the scatter points corresponds to H3K4me3 enrichment categories determined by density cutoffs (grey dotted lines, see Fig. S6D-G): yellow points = H3K4me3 enrichment in both sperm and pre-implantation embryos; light green points = H3K4me3 enrichment in only pre-implantation embryos; dark green points = H3K4me3 in sperm; blue points = absence of H3K4me3 enrichment in sperm and pre-implantation embryos. Represented pre-implantation embryo stages are 4-cell embryos (A), 8-cell embryos (B), and ICM (C). (D - G) Distribution of log2 (H3K4me3 counts + 1) at +/- 1 kb TSS of the hg19 genome in sperm (D), 4-cell embryos (E), 8-cell embryos (F), ICM (G). The local minimum was identified on the density plots (brown line) and used as the cutoff threshold value to identify promoters enriched for H3K4me3 in sperm and pre-implantation embryos (see Fig. S6A-C). (H - J) Scatterplots where the x axis corresponds to the log2 (pre-implantation embryo H3K4me3 promoter counts + 1) and the y axis corresponds to the log2 (sperm H3K4me3 promoter counts + 1) at +/- 1 kb TSS of the hg19 genome. Color of the scatter points correspond to the log2 pre-implantation embryo RPKM gene expression + 1. Dashed lines correspond to H3K4me3 promoter density cutoffs for pre-implantation embryo (x axis) or sperm (y axis). Represented pre-implantation embryo stages are 4-cell embryos (H), 8-cell embryos (I), and ICM (J). (K - M) Selected significant pathways from gene ontology analysis on promoters with H3K4me3 loss in South African sperm that retain H3K4me3 in the pre-implantation embryo and are expressed at the associated pre-implantation embryo stage (RPKM > 1; weighed Fisher p < 0.05). Size of dots corresponds to the number of genes from a significant pathway that overlap a peak with H3K4me3 gain. Color of the dots indicates -log2(weightFisher) value of significant pathway. Represented pre-implantation embryo stages are 4-cell embryos (K), 8-cell embryos (L), and ICM (M).

## References

1. Akbarian, S., Smith, M.A., and Jones, E.G. (1995). Editing for an AMPA receptor subunit RNA in prefrontal cortex and striatum in Alzheimer’s disease, Huntington’s disease and schizophrenia. Brain Research 699, 297–304. https://doi.org/10.1016/0006-8993(95)00922-D.

2. Alan S. Brown, M.D., M.P.H., Keely Cheslack-Postava, Ph.D., Panu Rantakokko, Ph.D., Hannu Kiviranta, Ph.D., Susanna Hinkka-Yli-Salomäki, Ph.D., Ian W. McKeague, Ph.D., Heljä-Marja Surcel, Ph.D., and Andre Sourander, M.D., Ph.D. (2018). Association of Maternal Insecticide Levels With Autism in Offspring From a National Birth Cohort. American Journal of Psychiatry 175, 1094–1101. 10.1176/appi.ajp.2018.17101129.

3. Alexa, A., Rahnenführer, J., and Lengauer, T. (2006). Improved scoring of functional groups from gene expression data by decorrelating GO graph structure. Bioinformatics 22, 1600–1607. 10.1093/bioinformatics/btl140.

4. Amemiya, H.M., Kundaje, A., and Boyle, A.P. (2019). The ENCODE Blacklist: Identification of Problematic Regions of the Genome. Scientific Reports 9, 9354. 10.1038/s41598-019-45839-z.

5. Aneck-Hahn, N.H., Schulenburg, G.W., Bornman, M.S., Farias, P., and de Jager, C. (2007). Impaired semen quality associated with environmental DDT exposure in young men living in a malaria area in the Limpopo Province, South Africa. J Androl 28, 423–434. 10.2164/jandrol.106.001701.

6. Anway, M.D., and Skinner, M.K. (2006). Epigenetic transgenerational actions of endocrine disruptors. Endocrinology 147, S43–49. 10.1210/en.2005-1058.

7. Aoshima, K., Inoue, E., Sawa, H., and Okada, Y. (2015). Paternal H3K4 methylation is required for minor zygotic gene activation and early mouse embryonic development. EMBO reports 16, 803–812.

8. Aston, K.I., Uren, P.J., Jenkins, T.G., Horsager, A., Cairns, B.R., Smith, A.D., and Carrell, D.T. (2015). Aberrant sperm DNA methylation predicts male fertility status and embryo quality. Fertility and Sterility 104, 1388–1397.e1385. 10.1016/j.fertnstert.2015.08.019.

9. Barakat, T.S., Halbritter, F., Zhang, M., Rendeiro, A.F., Perenthaler, E., Bock, C., and Chambers, I. (2018). Functional Dissection of the Enhancer Repertoire in Human Embryonic Stem Cells. Cell Stem Cell 23, 276–288.e278. https://doi.org/10.1016/j.stem.2018.06.014.

10. Benjamini, Y., and Hochberg, Y. (1995). Controlling the False Discovery Rate: A Practical and Powerful Approach to Multiple Testing. Journal of the Royal Statistical Society. Series B (Methodological) 57, 289–300.

11. Bestor, T.H. (1998). Cytosine methylation and the unequal developmental potentials of the oocyte and sperm genomes. The American Journal of Human Genetics 62, 1269–1273.

12. Bitman, J., Cecil, H.C., Harris, S.J., and Fries, G.F. (1968). Estrogenic Activity of o,p’-DDT in the Mammalian Uterus and Avian Oviduct. Science 162, 371–372. doi:10.1126/science.162.3851.371.

13. Bolger, A.M., Lohse, M., and Usadel, B. (2014). Trimmomatic: a flexible trimmer for Illumina sequence data. Bioinformatics (Oxford, England) 30, 2114–2120. 10.1093/bioinformatics/btu170.

14. Bonde, J.P., Toft, G., Rylander, L., Rignell-Hydbom, A., Giwercman, A., Spano, M., Manicardi, G.C., Bizzaro, D., Ludwicki, J.K., Zvyezday, V., et al. (2008). Fertility and markers of male reproductive function in Inuit and European populations spanning large contrasts in blood levels of persistent organochlorines. Environ Health Perspect 116, 269–277. 10.1289/ehp.10700.

15. Bornman, M.S., Chevrier, J., Rauch, S., Crause, M., Obida, M., Sathyanarayana, S., Barr, D.B., and Eskenazi, B. (2016). Dichlorodiphenyltrichloroethane exposure and anogenital distance in the Venda Health Examination of Mothers, Babies and their Environment (VHEMBE) birth cohort study, South Africa. Andrology 4, 608–615. 10.1111/andr.12235.

16. Bornman, R., De Jager, C., Worku, Z., Farias, P., and Reif, S. (2010). DDT and urogenital malformations in newborn boys in a malarial area. BJU International 106, 405–411. https://doi.org/10.1111/j.1464-410X.2009.09003.x.

17. Boucher, O., Jacobson Sandra, W., Plusquellec, P., Dewailly, É., Ayotte, P., Forget-Dubois, N., Jacobson Joseph, L., and Muckle, G. (2012). Prenatal Methylmercury, Postnatal Lead Exposure, and Evidence of Attention Deficit/Hyperactivity Disorder among Inuit Children in Arctic Québec. Environmental Health Perspectives 120, 1456–1461. 10.1289/ehp.1204976.

18. Bourc’his, D., and Bestor, T.H. (2004). Meiotic catastrophe and retrotransposon reactivation in male germ cells lacking Dnmt3L. Nature 431, 96–99.

19. Bourgey, M., Dali, R., Eveleigh, R., Chen, K.C., Letourneau, L., Fillon, J., Michaud, M., Caron, M., Sandoval, J., Lefebvre, F., et al. (2019). GenPipes: an open-source framework for distributed and scalable genomic analyses. Gigascience 8, giz037. 10.1093/gigascience/giz037.

20. Bouwman, H., and Kylin, H. (2009). Malaria control insecticide residues in breast milk: the need to consider infant health risks. Environmental health perspectives 117, 1477–1480. 10.1289/ehp.0900605.

21. Brown, T.M., Macdonald, R.W., Muir, D.C.G., and Letcher, R.J. (2018). The distribution and trends of persistent organic pollutants and mercury in marine mammals from Canada’s Eastern Arctic. Science of The Total Environment 618, 500–517. https://doi.org/10.1016/j.scitotenv.2017.11.052.

22. Brykczynska, U., Hisano, M., Erkek, S., Ramos, L., Oakeley, E.J., Roloff, T.C., Beisel, C., Schübeler, D., Stadler, M.B., and Peters, A.H. (2010). Repressive and active histone methylation mark distinct promoters in human and mouse spermatozoa. Nature structural & molecular biology 17, 679–687.

23. Cao, M., Shao, X., Chan, P., Cheung, W., Kwan, T., Pastinen, T., and Robaire, B. (2020). High-resolution analyses of human sperm dynamic methylome reveal thousands of novel age-related epigenetic alterations. Clinical Epigenetics 12, 192. 10.1186/s13148-020-00988-1.

24. Caporale, N., Leemans, M., Birgersson, L., Germain, P.L., Cheroni, C., Borbély, G., Engdahl, E., Lindh, C., Bressan, R.B., Cavallo, F., et al. (2022). From cohorts to molecules: Adverse impacts of endocrine disrupting mixtures. Science 375, eabe8244. 10.1126/science.abe8244.

25. Cartier, C., Muckle, G., Jacobson, S.W., Jacobson, J.L., Dewailly, E., Ayotte, P., Chevrier, C., and Saint-Amour, D. (2014). Prenatal and 5-year *p,p’*-DDE exposures are associated with altered sensory processing in school-aged children in Nunavik: a visual evoked potential study. Neurotoxicology 44, 8–16. 10.1016/j.neuro.2014.04.009.

26. Chan, D., Shao, X., Dumargne, M.-C., Aarabi, M., Simon, M.-M., Kwan, T., Bailey, J.L., Robaire, B., Kimmins, S., Gabriel, M.C.S., et al. (2019). Customized MethylC-Capture Sequencing to Evaluate Variation in the Human Sperm DNA Methylome Representative of Altered Folate Metabolism. Environmental Health Perspectives 127, 087002. doi:10.1289/EHP4812.

27. Chen, Q., Yan, M., Cao, Z., Li, X., Zhang, Y., Shi, J., Feng, G.H., Peng, H., Zhang, X., Zhang, Y., et al. (2016). Sperm tsRNAs contribute to intergenerational inheritance of an acquired metabolic disorder. Science 351, 397–400. 10.1126/science.aad7977.

28. Cheroni, C., Caporale, N., and Testa, G. (2020). Autism spectrum disorder at the crossroad between genes and environment: contributions, convergences, and interactions in ASD developmental pathophysiology. Molecular Autism 11, 69. 10.1186/s13229-020-00370-1.

29. Chuong, E.B., Rumi, M.A.K., Soares, M.J., and Baker, J.C. (2013). Endogenous retroviruses function as species-specific enhancer elements in the placenta. Nature Genetics 45, 325–329. 10.1038/ng.2553.

30. Consales, C., Toft, G., Leter, G., Bonde, J.P., Uccelli, R., Pacchierotti, F., Eleuteri, P., Jönsson, B.A., Giwercman, A., Pedersen, H.S., et al. (2016). Exposure to persistent organic pollutants and sperm DNA methylation changes in Arctic and European populations. Environ Mol Mutagen 57, 200–209. 10.1002/em.21994.

31. Corbin, J.M., and Ruiz-Echevarría, M.J. (2016). One-Carbon Metabolism in Prostate Cancer: The Role of Androgen Signaling. Int J Mol Sci 17. 10.3390/ijms17081208.

32. de Jager, C., Aneck-Hahn, N.H., Bornman, M.S., Farias, P., Leter, G., Eleuteri, P., Rescia, M., and Spanò, M. (2009). Sperm chromatin integrity in DDT-exposed young men living in a malaria area in the Limpopo Province, South Africa. Human Reproduction 24, 2429–2438. 10.1093/humrep/dep249.

33. Després, C., Beuter, A., Richer, F., Poitras, K., Veilleux, A., Ayotte, P., Dewailly, É., Saint-Amour, D., and Muckle, G. (2005). Neuromotor functions in Inuit preschool children exposed to Pb, PCBs, and Hg. Neurotoxicology and Teratology 27, 245–257. https://doi.org/10.1016/j.ntt.2004.12.001.

34. Diehl, A.G., Ouyang, N., and Boyle, A.P. (2020). Transposable elements contribute to cell and species-specific chromatin looping and gene regulation in mammalian genomes. Nature Communications 11, 1796. 10.1038/s41467-020-15520-5.

35. Ding, R., Jin, Y., Liu, X., Ye, H., Zhu, Z., Zhang, Y., Wang, T., and Xu, Y. (2017). Dose- and time- effect responses of DNA methylation and histone H3K9 acetylation changes induced by traffic-related air pollution. Scientific Reports 7, 43737. 10.1038/srep43737.

36. Dolinoy, D.C., Huang, D., and Jirtle, R.L. (2007). Maternal nutrient supplementation counteracts bisphenol A-induced DNA hypomethylation in early development. Proc Natl Acad Sci U S A 104, 13056–13061. 10.1073/pnas.0703739104.

37. Donkin, I., Versteyhe, S., Ingerslev, Lars R., Qian, K., Mechta, M., Nordkap, L., Mortensen, B., Appel, Emil Vincent R., Jørgensen, N., Kristiansen, Viggo B., et al. (2016). Obesity and Bariatric Surgery Drive Epigenetic Variation of Spermatozoa in Humans. Cell Metabolism 23, 369–378. https://doi.org/10.1016/j.cmet.2015.11.004.

38. Erkek, S., Hisano, M., Liang, C.-Y., Gill, M., Murr, R., Dieker, J., Schübeler, D., Vlag, J.v.d., Stadler, M.B., and Peters, A.H. (2013). Molecular determinants of nucleosome retention at CpG-rich sequences in mouse spermatozoa. Nature structural & molecular biology 20, 868–875.

39. Eskenazi, B., An, S., Rauch Stephen, A., Coker Eric, S., Maphula, A., Obida, M., Crause, M., Kogut Katherine, R., Bornman, R., and Chevrier, J. (2018). Prenatal Exposure to DDT and Pyrethroids for Malaria Control and Child Neurodevelopment: The VHEMBE Cohort, South Africa. Environmental Health Perspectives 126, 047004. 10.1289/EHP2129.

40. Eskenazi, B., Chevrier, J., Rosas, L.G., Anderson, H.A., Bornman, M.S., Bouwman, H., Chen, A., Cohn, B.A., de Jager, C., Henshel, D.S., et al. (2009). The Pine River statement: human health consequences of DDT use. Environmental health perspectives 117, 1359–1367. 10.1289/ehp.11748.

41. Feinberg, J.I., Bakulski, K.M., Jaffe, A.E., Tryggvadottir, R., Brown, S.C., Goldman, L.R., Croen, L.A., Hertz-Picciotto, I., Newschaffer, C.J., Daniele Fallin, M., and Feinberg, A.P. (2015). Paternal sperm DNA methylation associated with early signs of autism risk in an autism-enriched cohort. International Journal of Epidemiology 44, 1199–1210. 10.1093/ije/dyv028.

42. Freeberg, M.A., Fromont, L.A., D’Altri, T., Romero, A.F., Ciges, Jorge I., Jene, A., Kerry, G., Moldes, M., Ariosa, R., Bahena, S., et al. (2022). The European Genome-phenome Archive in 2021. Nucleic Acids Research 50, D980–D987. 10.1093/nar/gkab1059.

43. Gamallat, Y., Fang, X., Mai, H., Liu, X., Li, H., Zhou, P., Han, D., Zheng, S., Liao, C., Yang, M., et al. (2021). Bi-allelic mutation in Fsip1 impairs acrosome vesicle formation and attenuates flagellogenesis in mice. Redox Biology 43, 101969. https://doi.org/10.1016/j.redox.2021.101969.

44. Gao, T., and Qian, J. (2020). EnhancerAtlas 2.0: an updated resource with enhancer annotation in 586 tissue/cell types across nine species. Nucleic Acids Research 48, D58–D64. 10.1093/nar/gkz980.

45. Gapp, K., van Steenwyk, G., Germain, P.L., Matsushima, W., Rudolph, K.L.M., Manuella, F., Roszkowski, M., Vernaz, G., Ghosh, T., Pelczar, P., et al. (2020). Alterations in sperm long RNA contribute to the epigenetic inheritance of the effects of postnatal trauma. Molecular Psychiatry 25, 2162–2174. 10.1038/s41380-018-0271-6.

46. Gel, B., Díez-Villanueva, A., Serra, E., Buschbeck, M., Peinado, M.A., and Malinverni, R. (2016). regioneR: an R/Bioconductor package for the association analysis of genomic regions based on permutation tests. Bioinformatics 32, 289–291. 10.1093/bioinformatics/btv562.

47. Göke, J., Lu, X., Chan, Y.S., Ng, H.H., Ly, L.H., Sachs, F., and Szczerbinska, I. (2015). Dynamic transcription of distinct classes of endogenous retroviral elements marks specific populations of early human embryonic cells. Cell Stem Cell 16, 135–141. 10.1016/j.stem.2015.01.005.

48. Goodier, J.L. (2016). Restricting retrotransposons: a review. Mob DNA 7, 16. 10.1186/s13100-016-0070-z.

49. Guo, H., Zhu, P., Yan, L., Li, R., Hu, B., Lian, Y., Yan, J., Ren, X., Lin, S., Li, J., et al. (2014). The DNA methylation landscape of human early embryos. Nature 511, 606–610. 10.1038/nature13544.

50. Hammoud, S.S., Nix, D.A., Zhang, H., Purwar, J., Carrell, D.T., and Cairns, B.R. (2009). Distinctive chromatin in human sperm packages genes for embryo development. Nature 460, 473–478. 10.1038/nature08162.

51. Harutyunyan, A.S., Krug, B., Chen, H., Papillon-Cavanagh, S., Zeinieh, M., De Jay, N., Deshmukh, S., Chen, C.C.L., Belle, J., Mikael, L.G., et al. (2019). H3K27M induces defective chromatin spread of PRC2-mediated repressive H3K27me2/me3 and is essential for glioma tumorigenesis. Nature Communications 10, 1262. 10.1038/s41467-019-09140-x.

52. Heard, E., and Martienssen, R.A. (2014). Transgenerational epigenetic inheritance: myths and mechanisms. Cell 157, 95–109. 10.1016/j.cell.2014.02.045.

53. Herst, P.M., Dalvai, M., Lessard, M., Charest, P.L., Navarro, P., Joly-Beauparlant, C., Droit, A., Trasler, J.M., Kimmins, S., MacFarlane, A.J., et al. (2019). Folic acid supplementation reduces multigenerational sperm miRNA perturbation induced by in utero environmental contaminant exposure. Environmental Epigenetics 5, dvz024. 10.1093/eep/dvz024.

54. Himeidan, Y.E., Hamid, E.E., Thalib, L., El Bashir, M.I., and Adam, I. (2007). Climatic variables and transmission of falciparum malaria in New Halfa, eastern Sudan.

55. Hisano, M., Erkek, S., Dessus-Babus, S., Ramos, L., Stadler, M.B., and Peters, A.H. (2013). Genome-wide chromatin analysis in mature mouse and human spermatozoa. Nat Protoc 8, 2449–2470. 10.1038/nprot.2013.145.

56. Ishiuchi, T., Abe, S., Inoue, K., Yeung, W.K.A., Miki, Y., Ogura, A., and Sasaki, H. (2021). Reprogramming of the histone H3. 3 landscape in the early mouse embryo. Nature structural & molecular biology 28, 38–49.

57. Ito, J., Sugimoto, R., Nakaoka, H., Yamada, S., Kimura, T., Hayano, T., and Inoue, I. (2017). Systematic identification and characterization of regulatory elements derived from human endogenous retroviruses. PLOS Genetics 13, e1006883. 10.1371/journal.pgen.1006883.

58. Jacques, P., Jeyakani, J., and Bourque, G. (2013). The majority of primate-specific regulatory sequences are derived from transposable elements. PLoS Genet 9, e1003504. 10.1371/journal.pgen.1003504.

59. Jönsson, B.A.G., Rylander, L., Lindh, C., Rignell-Hydbom, A., Giwercman, A., Toft, G., Pedersen, H.S., Ludwicki, J.K., Góralczyk, K., Zvyezday, V., et al. (2005). Inter-population variations in concentrations, determinants of and correlations between 2,2’,4,4’,5,5’-hexachlorobiphenyl (CB-153) and 1,1-dichloro-2,2-bis (p-chlorophenyl)-ethylene (*p,p’*-DDE): a cross-sectional study of 3161 men and women from Inuit and European populations. Environmental Health 4, 27. 10.1186/1476-069X-4-27.

60. Kelce, W.R., Stone, C.R., Laws, S.C., Gray, L.E., Kemppainen, J.A., and Wilson, E.M. (1995). Persistent DDT metabolite *p,p’*-DDE is a potent androgen receptor antagonist. Nature 375, 581–585. 10.1038/375581a0.

61. Kimmins, S., and Sassone-Corsi, P. (2005). Chromatin remodelling and epigenetic features of germ cells. Nature 434, 583–589. 10.1038/nature03368.

62. King, S.E., McBirney, M., Beck, D., Sadler-Riggleman, I., Nilsson, E., and Skinner, M.K. (2019). Sperm epimutation biomarkers of obesity and pathologies following DDT induced epigenetic transgenerational inheritance of disease. Environmental Epigenetics 5, dvz008. 10.1093/eep/dvz008.

63. Krueger, F. (2015). Trim galore. A wrapper tool around Cutadapt and FastQC to consistently apply quality and adapter trimming to FastQ files 516, 517.

64. Krueger, F., and Andrews, S.R. (2011). Bismark: a flexible aligner and methylation caller for Bisulfite-Seq applications. Bioinformatics 27, 1571–1572. 10.1093/bioinformatics/btr167.

65. Küry, P., Nath, A., Créange, A., Dolei, A., Marche, P., Gold, J., Giovannoni, G., Hartung, H.P., and Perron, H. (2018). Human Endogenous Retroviruses in Neurological Diseases. Trends Mol Med 24, 379–394. 10.1016/j.molmed.2018.02.007.

66. Lambrot, R., Chan, D., Shao, X., Aarabi, M., Kwan, T., Bourque, G., Moskovtsev, S., Librach, C., Trasler, J., Dumeaux, V., and Kimmins, S. (2021). Whole-genome sequencing of H3K4me3 and DNA methylation in human sperm reveals regions of overlap linked to fertility and development. Cell Reports 36, 109418. https://doi.org/10.1016/j.celrep.2021.109418.

67. Lambrot, R., Siklenka, K., Lafleur, C., and Kimmins, S. (2019). The genomic distribution of histone H3K4me2 in spermatogonia is highly conserved in sperm†. Biology of Reproduction 100, 1661–1672. 10.1093/biolre/ioz055.

68. Lambrot, R., Xu, C., Saint-Phar, S., Chountalos, G., Cohen, T., Paquet, M., Suderman, M., Hallett, M., and Kimmins, S. (2013). Low paternal dietary folate alters the mouse sperm epigenome and is associated with negative pregnancy outcomes. Nature Communications 4, 2889. 10.1038/ncomms3889.

69. Lanciano, S., and Cristofari, G. (2020). Measuring and interpreting transposable element expression. Nature Reviews Genetics 21, 721–736. 10.1038/s41576-020-0251-y.

70. Langmead, B., and Salzberg, S.L. (2012). Fast gapped-read alignment with Bowtie 2. Nature Methods 9, 357–359. 10.1038/nmeth.1923.

71. Larose, H., Shami, A.N., Abbott, H., Manske, G., Lei, L., and Hammoud, S.S. (2019). Chapter Eight - Gametogenesis: A journey from inception to conception. In Current Topics in Developmental Biology, D.M. Wellik, ed. (Academic Press), pp. 257–310. https://doi.org/10.1016/bs.ctdb.2018.12.006.

72. Lesch, B.J., Silber, S.J., McCarrey, J.R., and Page, D.C. (2016). Parallel evolution of male germline epigenetic poising and somatic development in animals. Nature genetics 48, 888–894.

73. Lesch, B.J., Tothova, Z., Morgan, E.A., Liao, Z., Bronson, R.T., Ebert, B.L., and Page, D.C. (2019). Intergenerational epigenetic inheritance of cancer susceptibility in mammals. Elife 8, e39380.

74. Lessard, M., Herst, P.M., Charest, P.L., Navarro, P., Joly-Beauparlant, C., Droit, A., Kimmins, S., Trasler, J., Benoit-Biancamano, M.-O., MacFarlane, A.J., et al. (2019). Prenatal Exposure to Environmentally-Relevant Contaminants Perturbs Male Reproductive Parameters Across Multiple Generations that are Partially Protected by Folic Acid Supplementation. Scientific Reports 9, 13829. 10.1038/s41598-019-50060-z.

75. Lismer, A., Dumeaux, V., Lafleur, C., Lambrot, R., Brind’Amour, J., Lorincz, M.C., and Kimmins, S. (2021). Histone H3 lysine 4 trimethylation in sperm is transmitted to the embryo and associated with diet-induced phenotypes in the offspring. Dev Cell 56, 671–686.e676. 10.1016/j.devcel.2021.01.014.

76. Lismer, A., Siklenka, K., Lafleur, C., Dumeaux, V., and Kimmins, S. (2020). Sperm histone H3 lysine 4 trimethylation is altered in a genetic mouse model of transgenerational epigenetic inheritance. Nucleic Acids Research 48, 11380–11393. 10.1093/nar/gkaa712.

77. Liu, S., Brind’Amour, J., Karimi, M.M., Shirane, K., Bogutz, A., Lefebvre, L., Sasaki, H., Shinkai, Y., and Lorincz, M.C. (2014). Setdb1 is required for germline development and silencing of H3K9me3-marked endogenous retroviruses in primordial germ cells. Genes Dev 28, 2041–2055. 10.1101/gad.244848.114.

78. Liu, Y., Siegmund, K.D., Laird, P.W., and Berman, B.P. (2012). Bis-SNP: combined DNA methylation and SNP calling for Bisulfite-seq data. Genome Biol 13, R61. 10.1186/gb-2012-13-7-r61.

79. Longnecker, M.P., Klebanoff, M.A., Zhou, H., and Brock, J.W. (2001). Association between maternal serum concentration of the DDT metabolite DDE and preterm and small-for-gestational-age babies at birth. The Lancet 358, 110–114. https://doi.org/10.1016/S0140-6736(01)05329-6.

80. Lun, A.T., and Smyth, G.K. (2016). csaw: a Bioconductor package for differential binding analysis of ChIP-seq data using sliding windows. Nucleic Acids Res 44, e45. 10.1093/nar/gkv1191.

81. Ly, L., Chan, D., Aarabi, M., Landry, M., Behan, N.A., MacFarlane, A.J., and Trasler, J. (2017). Intergenerational impact of paternal lifetime exposures to both folic acid deficiency and supplementation on reproductive outcomes and imprinted gene methylation. Mol Hum Reprod 23, 461–477. 10.1093/molehr/gax029.

82. Lynch, V.J., Leclerc, R.D., May, G., and Wagner, G.P. (2011). Transposon-mediated rewiring of gene regulatory networks contributed to the evolution of pregnancy in mammals. Nat Genet 43, 1154–1159. 10.1038/ng.917.

83. Lynch, V.J., Nnamani, M.C., Kapusta, A., Brayer, K., Plaza, S.L., Mazur, E.C., Emera, D., Sheikh, S.Z., Grützner, F., Bauersachs, S., et al. (2015). Ancient transposable elements transformed the uterine regulatory landscape and transcriptome during the evolution of mammalian pregnancy. Cell Rep 10, 551–561. 10.1016/j.celrep.2014.12.052.

84. Maezawa, S., Yukawa, M., Alavattam, K.G., Barski, A., and Namekawa, S.H. (2018). Dynamic reorganization of open chromatin underlies diverse transcriptomes during spermatogenesis. Nucleic Acids Res 46, 593–608. 10.1093/nar/gkx1052.

85. Maurice, C., Dalvai, M., Lambrot, R., Deschênes, A., Scott-Boyer, M.-P., McGraw, S., Chan, D., Côté, N., Ziv-Gal, A., Flaws, J.A., et al. (2021). Early-Life Exposure to Environmental Contaminants Perturbs the Sperm Epigenome and Induces Negative Pregnancy Outcomes for Three Generations via the Paternal Lineage. Epigenomes 5, 10.

86. Michealraj, K.A., Kumar, S.A., Kim, L.J.Y., Cavalli, F.M.G., Przelicki, D., Wojcik, J.B., Delaidelli, A., Bajic, A., Saulnier, O., MacLeod, G., et al. (2020). Metabolic Regulation of the Epigenome Drives Lethal Infantile Ependymoma. Cell 181, 1329–1345.e1324. 10.1016/j.cell.2020.04.047.

87. Mochida, G.H., Mahajnah, M., Hill, A.D., Basel-Vanagaite, L., Gleason, D., Hill, R.S., Bodell, A., Crosier, M., Straussberg, R., and Walsh, C.A. (2009). A Truncating Mutation of TRAPPC9 Is Associated with Autosomal-Recessive Intellectual Disability and Postnatal Microcephaly. The American Journal of Human Genetics 85, 897–902. https://doi.org/10.1016/j.ajhg.2009.10.027.

88. Molaro, A., Hodges, E., Fang, F., Song, Q., McCombie, W.R., Hannon, Gregory J., and Smith, Andrew D. (2011). Sperm Methylation Profiles Reveal Features of Epigenetic Inheritance and Evolution in Primates. Cell 146, 1029–1041. https://doi.org/10.1016/j.cell.2011.08.016.

89. Muir, D.C.G., Wagemann, R., Hargrave, B.T., Thomas, D.J., Peakall, D.B., and Norstrom, R.J. (1992). Arctic marine ecosystem contamination. Science of The Total Environment 122, 75–134. https://doi.org/10.1016/0048-9697(92)90246-O.

90. Niedzwiecki Megan, M., Hall Megan, N., Liu, X., Oka, J., Harper Kristin, N., Slavkovich, V., Ilievski, V., Levy, D., van Geen, A., Mey Jacob, L., et al. (2013). A Dose–Response Study of Arsenic Exposure and Global Methylation of Peripheral Blood Mononuclear Cell DNA in Bangladeshi Adults. Environmental Health Perspectives 121, 1306–1312. 10.1289/ehp.1206421.

91. Nishikura, K. (2010). Functions and regulation of RNA editing by ADAR deaminases. Annual review of biochemistry 79, 321–349.

92. Okano, M., Bell, D.W., Haber, D.A., and Li, E. (1999). DNA methyltransferases Dnmt3a and Dnmt3b are essential for de novo methylation and mammalian development. Cell 99, 247–257.

93. Oluwayiose, O.A., Marcho, C., Wu, H., Houle, E., Krawetz, S.A., Suvorov, A., Mager, J., and Richard Pilsner, J. (2021). Paternal preconception phthalate exposure alters sperm methylome and embryonic programming. Environ Int 155, 106693. 10.1016/j.envint.2021.106693.

94. Ooi, S.K., Qiu, C., Bernstein, E., Li, K., Jia, D., Yang, Z., Erdjument-Bromage, H., Tempst, P., Lin, S.-P., and Allis, C.D. (2007). DNMT3L connects unmethylated lysine 4 of histone H3 to de novo methylation of DNA. Nature 448, 714–717.

95. Organisation, W.H. (1999). WHO laboratory manual for the examination of human semen and sperm-cervical mucus interaction (Cambridge university press).

96. Patel, D.M., Jones, R.R., Booth, B.J., Olsson, A.C., Kromhout, H., Straif, K., Vermeulen, R., Tikellis, G., Paltiel, O., Golding, J., et al. (2020). Parental occupational exposure to pesticides, animals and organic dust and risk of childhood leukemia and central nervous system tumors: Findings from the International Childhood Cancer Cohort Consortium (I4C). Int J Cancer 146, 943–952. 10.1002/ijc.32388.

97. Paz, N., Levanon, E.Y., Amariglio, N., Heimberger, A.B., Ram, Z., Constantini, S., Barbash, Z.S., Adamsky, K., Safran, M., Hirschberg, A., et al. (2007). Altered adenosine-to-inosine RNA editing in human cancer. Genome Res 17, 1586–1595. 10.1101/gr.6493107.

98. Pepin, A.-S., Lafleur, C., Lambrot, R., Dumeaux, V., and Kimmins, S. (2022). Sperm Histone H3 Lysine 4 tri- methylation serves as a metabolic sensor of paternal obesity and is associated with the inheritance of metabolic dysfunction. Molecular Metabolism, 101463. https://doi.org/10.1016/j.molmet.2022.101463.

99. Radford, E.J., Ito, M., Shi, H., Corish, J.A., Yamazawa, K., Isganaitis, E., Seisenberger, S., Hore, T.A., Reik, W., Erkek, S., et al. (2014). In utero effects. In utero undernourishment perturbs the adult sperm methylome and intergenerational metabolism. Science 345, 1255903. 10.1126/science.1255903.

100. Reed, M.L., and Leff, S.E. (1994). Maternal imprinting of human SNRPN, a gene deleted in Prader–Willi syndrome. Nature Genetics 6, 163–167. 10.1038/ng0294-163.

101. Richthoff, J., Rylander, L., Jönsson, B.A., Akesson, H., Hagmar, L., Nilsson-Ehle, P., Stridsberg, M., and Giwercman, A. (2003). Serum levels of 2,2’,4,4’,5,5’-hexachlorobiphenyl (CB-153) in relation to markers of reproductive function in young males from the general Swedish population. Environ Health Perspect 111, 409–413. 10.1289/ehp.5767.

102. Rignell-Hydbom, A., Rylander, L., Giwercman, A., Jönsson, B.A., Nilsson-Ehle, P., and Hagmar, L. (2004). Exposure to CB-153 and *p,p’*-DDE and male reproductive function. Hum Reprod 19, 2066–2075. 10.1093/humrep/deh362.

103. Rodriguez-Terrones, D., and Torres-Padilla, M.E. (2018). Nimble and Ready to Mingle: Transposon Outbursts of Early Development. Trends Genet 34, 806–820. 10.1016/j.tig.2018.06.006.

104. Rosenquist Aske, H., Høyer Birgit, B., Julvez, J., Sunyer, J., Pedersen Henning, S., Lenters, V., Jönsson Bo, A.G., Bonde Jens, P., and Toft, G. (2017). Prenatal and Postnatal PCB-153 and *p,p′*-DDE Exposures and Behavior Scores at 5–9 Years of Age among Children in Greenland and Ukraine. Environmental Health Perspectives 125, 107002. 10.1289/EHP553.

105. Santenard, A., Ziegler-Birling, C., Koch, M., Tora, L., Bannister, A.J., and Torres-Padilla, M.-E. (2010). Heterochromatin formation in the mouse embryo requires critical residues of the histone variant H3. 3. Nature cell biology 12, 853–862.

106. Schmidt, D., Schwalie, P.C., Wilson, M.D., Ballester, B., Gonçalves, A., Kutter, C., Brown, G.D., Marshall, A., Flicek, P., and Odom, D.T. (2012). Waves of retrotransposon expansion remodel genome organization and CTCF binding in multiple mammalian lineages. Cell 148, 335–348. 10.1016/j.cell.2011.11.058.

107. Senft, A.D., and Macfarlan, T.S. (2021). Transposable elements shape the evolution of mammalian development. Nature Reviews Genetics 22, 691–711. 10.1038/s41576-021-00385-1.

108. Severinsen, J.E., Bjarkam, C.R., Kiar-Larsen, S., Olsen, I.M., Nielsen, M.M., Blechingberg, J., Nielsen, A.L., Holm, I.E., Foldager, L., Young, B.D., et al. (2006). Evidence implicating BRD1 with brain development and susceptibility to both schizophrenia and bipolar affective disorder. Molecular Psychiatry 11, 1126–1138. 10.1038/sj.mp.4001885.

109. Shabalin, A.A. (2012). Matrix eQTL: ultra fast eQTL analysis via large matrix operations. Bioinformatics 28, 1353–1358. 10.1093/bioinformatics/bts163.

110. Sharma, U., Conine, C.C., Shea, J.M., Boskovic, A., Derr, A.G., Bing, X.Y., Belleannee, C., Kucukural, A., Serra, R.W., Sun, F., et al. (2016). Biogenesis and function of tRNA fragments during sperm maturation and fertilization in mammals. Science 351, 391–396. 10.1126/science.aad6780.

111. Shim, Y.K., Mlynarek, S.P., and Wijngaarden, E.v. (2009). Parental Exposure to Pesticides and Childhood Brain Cancer: U.S. Atlantic Coast Childhood Brain Cancer Study. Environmental Health Perspectives 117, 1002–1006. doi:10.1289/ehp.0800209.

112. Siklenka, K., Erkek, S., Godmann, M., Lambrot, R., McGraw, S., Lafleur, C., Cohen, T., Xia, J., Suderman, M., Hallett, M., et al. (2015). Disruption of histone methylation in developing sperm impairs offspring health transgenerationally. Science 350, aab2006. 10.1126/science.aab2006.

113. Simonich Staci, L., and Hites Ronald, A. (1995). Global Distribution of Persistent Organochlorine Compounds. Science 269, 1851–1854. 10.1126/science.7569923.

114. Skinner, M.K., Ben Maamar, M., Sadler-Riggleman, I., Beck, D., Nilsson, E., McBirney, M., Klukovich, R., Xie, Y., Tang, C., and Yan, W. (2018). Alterations in sperm DNA methylation, non-coding RNA and histone retention associate with DDT-induced epigenetic transgenerational inheritance of disease. Epigenetics Chromatin 11, 8. 10.1186/s13072-018-0178-0.

115. Skinner, M.K., Manikkam, M., Tracey, R., Guerrero-Bosagna, C., Haque, M., and Nilsson, E.E. (2013). Ancestral dichlorodiphenyltrichloroethane (DDT) exposure promotes epigenetic transgenerational inheritance of obesity. BMC Medicine 11, 228. 10.1186/1741-7015-11-228.

116. Skvortsova, K., Iovino, N., and Bogdanović, O. (2018). Functions and mechanisms of epigenetic inheritance in animals. Nat Rev Mol Cell Biol 19, 774–790. 10.1038/s41580-018-0074-2.

117. Soubry, A., Hoyo, C., Jirtle, R.L., and Murphy, S.K. (2014). A paternal environmental legacy: evidence for epigenetic inheritance through the male germ line. Bioessays 36, 359–371. 10.1002/bies.201300113.

118. Spanò, M., Toft, G., Hagmar, L., Eleuteri, P., Rescia, M., Rignell-Hydbom, A., Tyrkiel, E., Zvyezday, V., and Bonde, J.P. (2005). Exposure to PCB and p, p’-DDE in European and Inuit populations: impact on human sperm chromatin integrity. Hum Reprod 20, 3488–3499. 10.1093/humrep/dei297.

119. Stringer, J.M., Forster, S.C., Qu, Z., Prokopuk, L., O’Bryan, M.K., Gardner, D.K., White, S.J., Adelson, D., and Western, P.S. (2018). Reduced PRC2 function alters male germline epigenetic programming and paternal inheritance. BMC biology 16, 1–20.

120. Tang, Walfred W.C., Dietmann, S., Irie, N., Leitch, Harry G., Floros, Vasileios I., Bradshaw, Charles R., Hackett, Jamie A., Chinnery, Patrick F., and Surani, M.A. (2015). A Unique Gene Regulatory Network Resets the Human Germline Epigenome for Development. Cell 161, 1453–1467. https://doi.org/10.1016/j.cell.2015.04.053.

121. Teran, T., Lamon, L., and Marcomini, A. (2012). Climate change effects on POPs’ environmental behaviour: a scientific perspective for future regulatory actions. Atmospheric Pollution Research 3, 466–476. https://doi.org/10.5094/APR.2012.054.

122. Toft, G., Axmon, A., Giwercman, A., Thulstrup, A.M., Rignell-Hydbom, A., Pedersen, H.S., Ludwicki, J.K., Zvyezday, V., Zinchuk, A., Spano, M., et al. (2005). Fertility in four regions spanning large contrasts in serum levels of widespread persistent organochlorines: a cross-sectional study. Environ Health 4, 26. 10.1186/1476-069x-4-26.

123. Toft, G., Rignell-Hydbom, A., Tyrkiel, E., Shvets, M., Giwercman, A., Lindh, C.H., Pedersen, H.S., Ludwicki, J.K., Lesovoy, V., Hagmar, L., et al. (2006). Semen Quality and Exposure to Persistent Organochlorine Pollutants. Epidemiology 17.

124. Turusov, V., Rakitsky, V., and Tomatis, L. (2002). Dichlorodiphenyltrichloroethane (DDT): ubiquity, persistence, and risks. Environmental Health Perspectives 110, 125–128. 10.1289/ehp.02110125.

125. van den Berg, H. (2010). DDT and Malaria Prevention: van den Berg Responds. Environmental Health Perspectives 118, A15–A16. 10.1289/ehp.0901276R.

126. van den Berg, H., Manuweera, G., and Konradsen, F. (2017). Global trends in the production and use of DDT for control of malaria and other vector-borne diseases. Malaria Journal 16, 401. 10.1186/s12936-017-2050-2.

127. Van Oostdam, J., Donaldson, S.G., Feeley, M., Arnold, D., Ayotte, P., Bondy, G., Chan, L., Dewaily, É., Furgal, C.M., Kuhnlein, H., et al. (2005). Human health implications of environmental contaminants in Arctic Canada: A review. Science of The Total Environment 351-352, 165–246. https://doi.org/10.1016/j.scitotenv.2005.03.034.

128. Veselovska, L., Smallwood, S.A., Saadeh, H., Stewart, K.R., Krueger, F., Maupetit-Méhouas, S., Arnaud, P., Tomizawa, S.-i., Andrews, S., and Kelsey, G. (2015). Deep sequencing and de novo assembly of the mouse oocyte transcriptome define the contribution of transcription to the DNA methylation landscape. Genome Biology 16, 209. 10.1186/s13059-015-0769-z.

129. Weihe, P., Debes, F., Halling, J., Petersen, M.S., Muckle, G., Odland, J.Ø., Dudarev, A.A., Ayotte, P., Dewailly, É., Grandjean, P., and Bonefeld-Jørgensen, E. (2016). Health effects associated with measured levels of contaminants in the Arctic. International Journal of Circumpolar Health 75, 33805. 10.3402/ijch.v75.33805.

130. World Health, O., United Nations Environment, P., Inter-Organization Programme for the Sound Management of, C., Bergman, Å., Heindel, J.J., Jobling, S., Kidd, K., and Zoeller, T.R. (2013). State of the science of endocrine disrupting chemicals 2012 : summary for decision-makers. World Health Organization. 2013. https://apps.who.int/iris/handle/10665/78102.

131. World Health Organization. Global Malaria, P., and World Health Organization. Malaria, U. (2006). Indoor residual spraying : use of indoor residual spraying for scaling up global malaria control and elimination : WHO position statement. World Health Organization.

132. Xia, W., Xu, J., Yu, G., Yao, G., Xu, K., Ma, X., Zhang, N., Liu, B., Li, T., Lin, Z., et al. (2019). Resetting histone modifications during human parental-to-zygotic transition. Science 365, 353–360. 10.1126/science.aaw5118.

133. Zeng, L., Wang, M., Zhou, J., Wang, X., Zhang, Y., and Su, P. (2022). A hypothesis: Retrotransposons as a relay of epigenetic marks in intergenerational epigenetic inheritance. Gene 817, 146229. 10.1016/j.gene.2022.146229.

134. Zhang, Y., Schöttker, B., Florath, I., Stock, C., Butterbach, K., Holleczek, B., Mons, U., and Brenner, H. (2016). Smoking-Associated DNA Methylation Biomarkers and Their Predictive Value for All-Cause and Cardiovascular Mortality. Environmental Health Perspectives 124, 67–74. 10.1289/ehp.1409020.

